# Broad de-regulated U2AF1 splicing is prognostic and augments leukemic transformation via protein arginine methyltransferase activation

**DOI:** 10.1101/2024.02.04.578798

**Authors:** Meenakshi Venkatasubramanian, Leya Schwartz, Nandini Ramachandra, Joshua Bennett, Krithika R. Subramanian, Xiaoting Chen, Shanisha Gordon-Mitchell, Ariel Fromowitz, Kith Pradhan, David Shechter, Srabani Sahu, Diane Heiser, Peggy Scherle, Kashish Chetal, Aishwarya Kulkarni, Kasiani C. Myers, Matthew T. Weirauch, H. Leighton Grimes, Daniel T. Starczynowski, Amit Verma, Nathan Salomonis

## Abstract

The role of splicing dysregulation in cancer is underscored by splicing factor mutations; however, its impact in the absence of such rare mutations is poorly understood. To reveal complex patient subtypes and putative regulators of pathogenic splicing in Acute Myeloid Leukemia (AML), we developed a new approach called OncoSplice. Among diverse new subtypes, OncoSplice identified a biphasic poor prognosis signature that partially phenocopies *U2AF1*-mutant splicing, impacting thousands of genes in over 40% of adult and pediatric AML cases. *U2AF1*-like splicing co-opted a healthy circadian splicing program, was stable over time and induced a leukemia stem cell (LSC) program. Pharmacological inhibition of the implicated *U2AF1*-like splicing regulator, PRMT5, rescued leukemia mis-splicing and inhibited leukemic cell growth. Genetic deletion of IRAK4, a common target of *U2AF1*-like and PRMT5 treated cells, blocked leukemia development in xenograft models and induced differentiation. These analyses reveal a new prognostic alternative-splicing mechanism in malignancy, independent of splicing-factor mutations.

**Statement of significance:** Using a new in silico strategy we reveal counteracting determinants of patient survival in Acute Myeloid Leukemia that co-opt well-defined mutation-dependent splicing programs. Broad poor-prognosis splicing and leukemia stem cell survival could be rescued through pharmacological inhibition (PRMT5) or target deletion (IRAK4), opening the door for new precision therapies.

**Competing Interests:** Conflict-of-interest disclosure: DTS. serves on the scientific advisory board at Kurome Therapeutics; is a consultant for and/or received funding from Kurome Therapeutics, Captor Therapeutics, Treeline Biosciences, and Tolero Therapeutics; and has equity in Kurome Therapeutics. AV has received research funding from GlaxoSmithKline, BMS, Jannsen, Incyte, MedPacto, Celgene, Novartis, Curis, Prelude and Eli Lilly and Company, has received compensation as a scientific advisor to Novartis, Stelexis Therapeutics, Acceleron Pharma, and Celgene, and has equity ownership in Throws Exception and Stelexis Therapeutics.

## INTRODUCTION

Alternative splicing is a primary mechanism used to achieve mRNA transcript and proteomic diversity in higher eukaryotes ^1^. In cancer, altered mRNA splicing can lead to aberrant protein products that promote oncogenic transformation and metastasis and confer chemotherapy resistance ^2–6^. In the absence of direct splicing mutations, other mechanisms exist to modify splicing pathways in diverse cancers including modulation of splicing factor gene expression and the mutation of splicing-factor-interacting proteins ^7–12^. One such example is the pathogenic splicing of the gene *IRAK4* in myelodysplastic syndromes (MDS) and AML, in which *IRAK4* exon-inclusion resulting in the expression of a hypermorphic IRAK4 isoform occurs in a large subset of these malignancies in the absence of its primary regulator (*U2AF1* mutations) ^13–15^. Thus, atypical, coordinated splicing may in general be a significant mediator of cancer and other complex diseases.

While alternative splicing is a recognized oncogenic driver in a small percentage of adult acute myeloid leukemia (AML) (∼10-15%), splicing factor mutations are rarely found in pediatric AML. These data suggest that pathogenic splicing is not a primary mediator of pediatric cancer survival or therapeutic response. Current approaches to assess the role of splicing in complex diseases rely on the focused analysis of prior-defined genetic, epigenetic or gene-expression subtypes ^16–19^. While in principle, the discovery of splicing-defined cancer subtypes should be comparable to gene expression, such analyses are hindered by the variable detection of isoform expression from RNA-Seq, complex overlapping genetics, tumor cellular heterogeneity and redundant events. Further, methods to predict likely causal splicing regulators from alternative splicing remain in their relative infancy, focused principally on the co-occurrence of alternative splicing with predicted cis-regulatory binding sites ^20,21^.

## RESULTS

### Unsupervised classification and regulatory prediction of cancer splicing subtypes

To characterize the splicing landscape of adult and pediatric AML we processed two available adult AML RNA-Seq datasets - Leucegene (437 adult) ^22^, and the Cancer Genome Atlas (TCGA, 179 adult) ^23^ and compared them to the pediatric AML dataset TARGET (390 pediatric patients with 257 at diagnosis) which lacks splicing-factor mutations ^24^. The genetics of the AML samples were determined from existing cancer databases and *de novo* RNA-Seq variant analysis (**ED Fig. 1a,b**). As expected, supervised analysis of cancer RNA-Seq data, identified distinct genomic lesions that result in highly-specific gene-expression and splicing signatures, including those in common adult and pediatric oncofusions (**Fig. 1a,b** and **ED Fig. 1c,d**). While such signatures enable highly accurate supervised classification, such subtypes could not be resolved by existing conventional unsupervised analyses (**Fig. 1c** and **ED Fig. 1e**). Such difficulty stems from overlapping signatures due to the presence of multiple genomic lesions per sample, as well as unknown splicing signatures that confound subtype identification.

**Figure 1.**
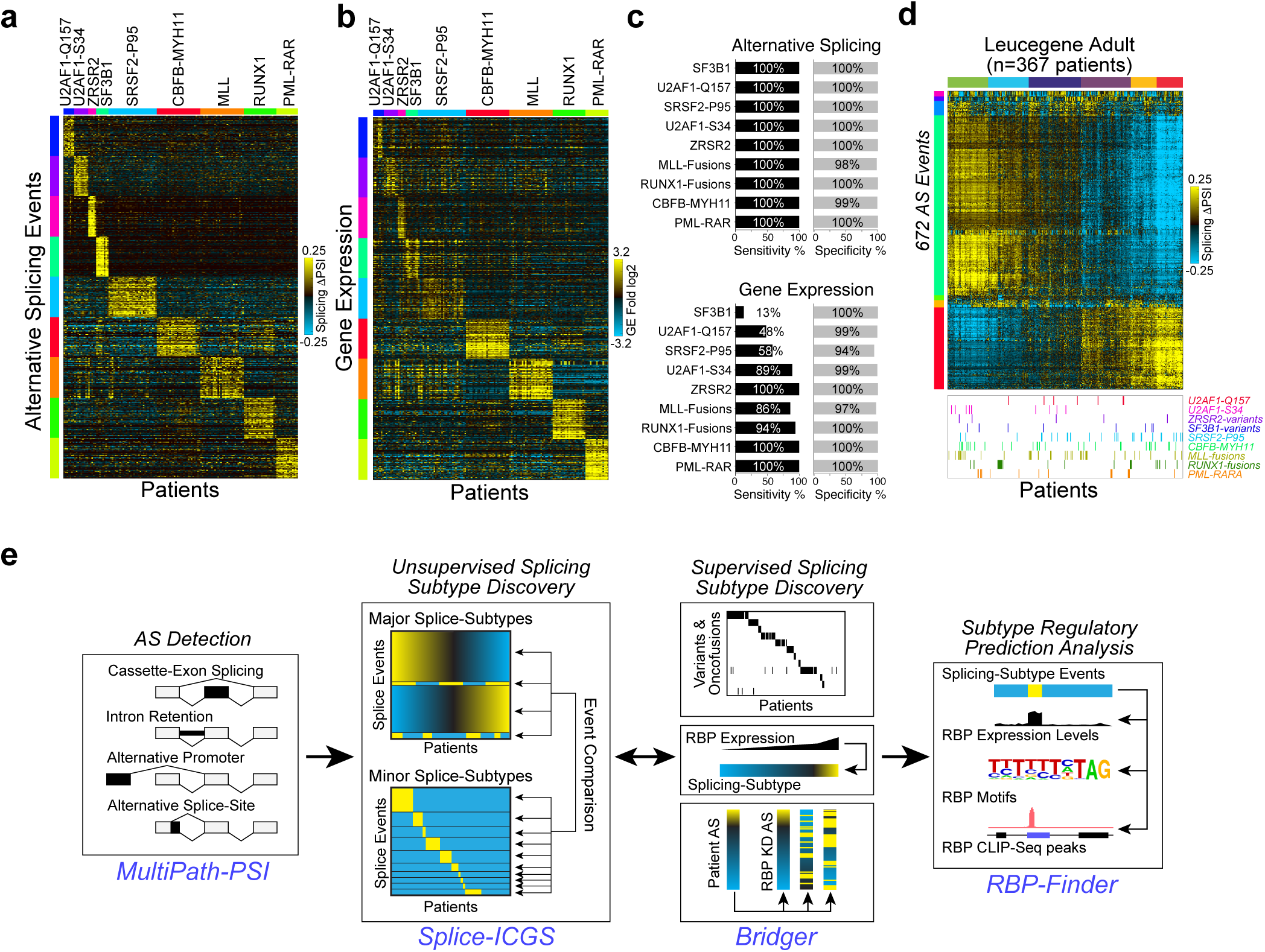
Mutation-defined splicing is largely obscured in leukemia. **a,b**) Heatmap of top marker splicing events (a) and differentially expressed genes (b) in AML (Leucegene RNA-Seq) for a subset of patients and common splicing factor mutations/fusions (n=142, traininig). c) Relative ability of splicing versus gene expression to accurately classify AML patient genetics (n=200), based on 3-fold cross-validation (SVM, one vs. rest). Columns=patients, Rows=events/genes. Delta PSI=relative difference in Percent Spliced In (PSI) values. d) Heatmap of alternative splicing-patterns identified in Leucegene RNA-Seq, identified using a single-cell analysis clustering algorithm (ICGS). a) Cartoon of the OncoSplice computational workflow to define new splicing subtypes and mechanisms of gene regulation from RNA-Seq. These steps consist of: 1) splicing quantification, 2) unsupervised subtype discovery, 3) supervised subtype identification (genetics, multi-factor splicing event correlation) and 4) RNA-regulatory splicing-subtype prediction based on RBP expression, binding sites and CLIP-Seq data.

To resolve complex overlapping splicing patterns and their mode of regulation, we developed a novel automated computational workflow termed OncoSplice **(Fig. 1e**). OncoSplice implements multiple algorithms, including: a) a highly accurate Percent-Spliced-In (ψ) algorithm (MultiPath-PSI) ^25^, b) unsupervised patient splicing-subtype detection (splice-ICGS), c) supervised mutation and subtype predictions (Bridger) and d) RNA-Binding Protein (RBP) regulatory prediction (RBP-Finder).

The principal innovation in OncoSplice is splice-ICGS, designed to identify novel patient subtypes defined uniquely by their splicing profiles. This workflow initially identifies variable coordinated splicing events using an adaptation of our single-cell RNA-Seq (scRNA-Seq) pipeline ICGS2 (Iterative Clustering and Guide-gene Selection) to enable multiclass assignments for all samples ^26^ (**ED Fig. 1f**). Because large-broad splicing patterns can confound the identification of rare splicing-defined patient subtypes (termed herein as splice-archetypes), splice-ICGS iterates its analysis, excluding all splicing events correlated to signatures identified in the prior round, to comprehensively define all splicing programs. This approach allows the same patients to occur in independent splice-archetypes, associated with distinct genetic drivers. Broad splicing signatures can represent novel splicing regulatory pathway differences, cell lineage differences or batch effects. When applied to a subset of AML patients with defined splicing factor mutations and oncofusions, splice-ICGS recovers nearly all known subtypes (8 out 9), whereas as cutting-edge scRNA-Seq or bi/fuzzy-clustering approaches capture only a few at most (**ED Fig. 1g**).

To identify additional splice-archetypes, OncoSplice finds genomic variants associated with distinct PSI programs, *de novo* PSI signatures correlated to RBP gene expression and associates tumor samples with specific disrupted RBPs based on their correlation to RBP-knockdown PSI profiles (Bridger module). Finally, the RBP-Finder module of OncoSplice associates regulatory RBP for each identified splice-archetype from PSI enrichment of RBP motifs, CLIP-Seq and RBP differential gene expression. This algorithm is a modification of our previous developed RELI algorithm for transcriptional regulation ^27^ in combination with a weighted logistic regression model (**Supplemental Methods**).

### OncoSplice identifies the spectrum of splicing-defined disease archetypes in AML

Application of OncoSplice to the majority of Leucegene AML samples identified a total of 25 splicing subtypes, of which 15 were previously defined in AML and 10 were novel subtypes (**Fig. 2a**, **ED Table 1**). The large majority of these subtypes (n=19) were specifically identified by splice-ICGS. One such subtype revealed *SRSF2* point mutations and in frame P95 to R102 8AA deletion ^28^ as a single related subtype . Only one of the OncoSplice subtypes was identified from comparison to RBP knockdown splicing profiles (Bridger) - 11 patients with confirmed heterogenous *HNRNPK* insertions/deletions, partial chromosomal deletions (partial 9q), or splicing defects in *HNRNPK* itself (intron retention). This *HNRNPK* subtype would not be identified through conventional genotype-based analyses, as it is produced by heterogenous genetic and non-genetic impacts. Frequently co-occurring variants among all subtypes were reported by OncoSplice, including enrichment of *KIT* mutations with *CBFB-MYH11*, *IDH2-R140Q* with *SRSF2-P95*, and *EVI* fusions with *SF3B1* variants. In addition, several of the most frequent and novel detected splicing-subtypes were enriched for known cancer variants, the most significant of which included *TP53* mutations (15%) or combined *FLT3* internal tandem duplications (FLT3-ITD) and NPM1 mutations (19%). Among these novel subtypes, two separate clusters were highly enriched for *TP53* mutations, with one of these specifically enriched for TCGA subtype annotations for acute erythroid leukemias (M6 subtype, R2-C3 cluster – **Table 1**). Likewise, two *NPM1-FLT3* enriched subtypes were found with coincident enrichment in either *TET2* (9%) or *DNMT3A* (10%). Among these mutation-enriched splicing subtypes, *NPM1-FLT3*-*DNMT3A* uniquely predicted poor overall survival in TCGA (**Table 1**). While such genetically defined subtypes have been previously defined ^29,30^, the unique combination of these mutations results in novel splicing subtypes that are distinct from each other and suggest a novel role for divergent epigenetic modulation of alternative splicing (*TET2* or *DNMT3A*).

**Figure 2.**
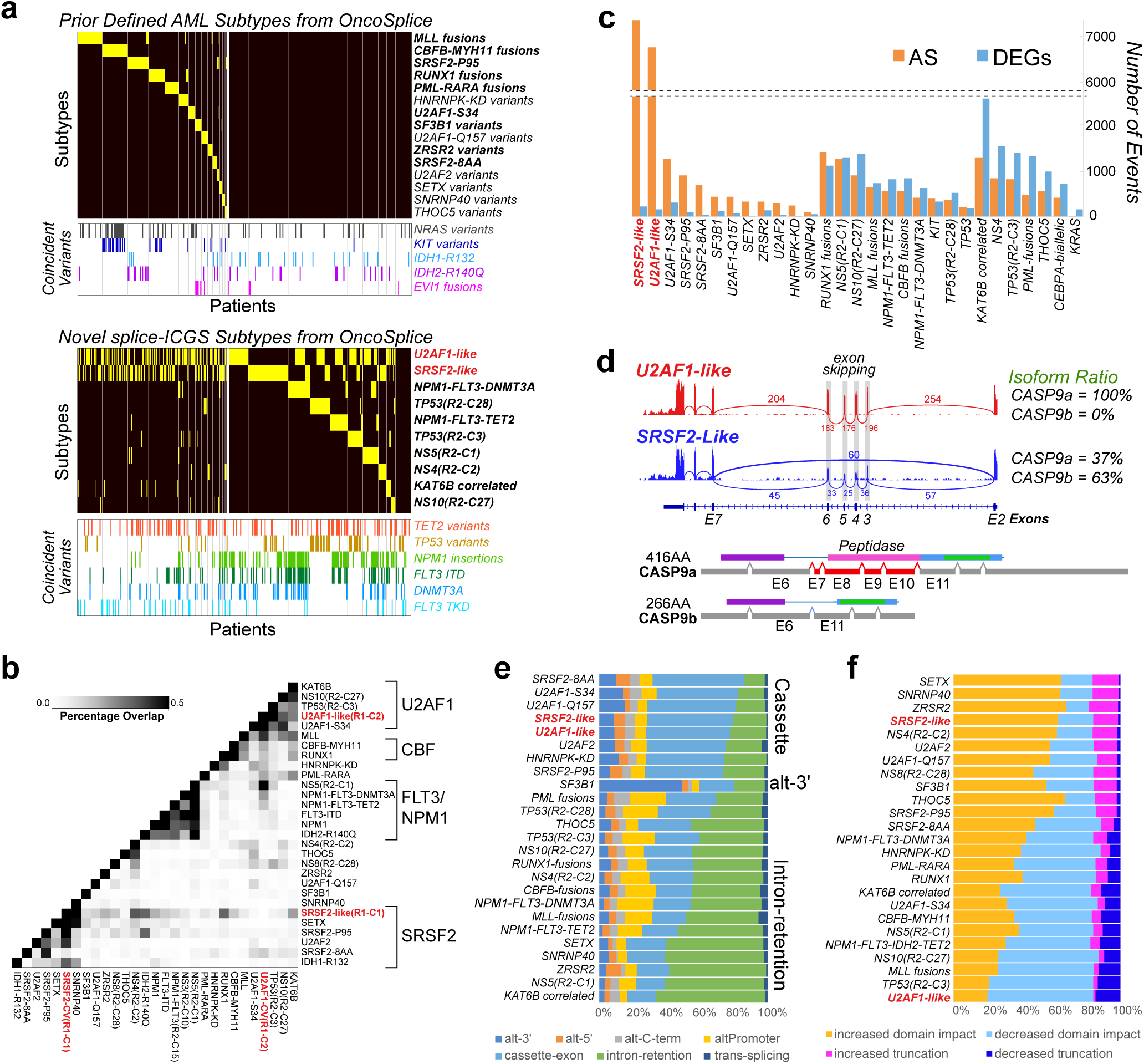
OncoSplice uncovers genetically heterogenous subtypes AML. **a**) OncoSplice-defined AML subtypes with coincident cancer genomic variants in 367 adult AML samples (yellow=subtype assigned patient) (ED Table 1). Previously-defined AML subtypes (top panel) and novel OncoSplice-defined subtypes (bottom panel), annotated for RNA-Seq-detected genomic variants, oncofusions, deletions or structural rearrangements (bold=splice-ICGS reported). For the top panel, final subtypes were revised according to known patient genetics (Supplementary Methods – Section 7). Note that *U2AF1*-like and *SRSF2*-like splicing subtypes co-occur with other splicing subtypes. **b**) Heatmap of concordant splicing events between OncoSplice-defined subtypes. Hierarchical clustering of the percentage of overlapping splicing events between all pairs of AML subtypes (regulated in the same direction) are shown (black=high percentage, white=low). Clusters of samples dominated by *U2AF1* (*U2AF1*-like or *U2AF1* mutation), *SRSF2* (*SRSF2-like* or *SRSF2* mutation) or NPM1/FLT3-ITD are labeled (right). c) For major OncoSplice-defined subtypes the number of differentially-expressed genes (DEGs) and unique alternative-splicing events (AS) are shown. Subtypes are grouped into those principally defined by AS (left), AS and DEGs (middle) or DEGs (right). The potentially confounding effect of *U2AF1*-like and *SRSF2*-like splicing events has been removed from the other subtypes. d) Splicing example: SashimiPlot of *CASP9* splicing in a *U2AF1*-like and an *SRSF2*-like patient sample (top). SashimiPlot lines between exons indicate junctions and numbers indicate junction-read counts. The alternative splice event results in predicted CASP9 protein isoforms (bottom) including the pro-apoptotic long CASP9a isoform and the short CASP9b isoform, which lacks the peptidase domain (ExonPlot view AltAnalyze). e) Annotation of the frequency of MultiPath-PSI-defined splice-event types (defined below) associated with each AML subtype (denoted to the left). f) Annotation of the AltAnalyze-predicted impact of splice events on protein domain and protein length in each AML subtype (denoted to the left).

**Table 1.** Broad characterization of novel adult and pediatric AML subtypes. All Leucegene identified splicing subtypes are shown with patient statistics for Leucegene and corresponding subtypes from TCGA and TARGET RNA-Seq (splice-ICGS supervised classification). Red = novel causal variants for known AML subtypes. RBP-Finder predicted regulatory RBPs are indicated for representative top scoring predictions. Asterisk = previously annotated cancer-associated splicing events. Survival is calculated based on the coxph test for each subtype and AML subtype associations for overall survival in patients under 60 years of age compared to cytogenetically normal AMLs. Blue = poor prognosis. Subtypes without a p-value are either not associated with survival (p<0.05) or have too few samples in TCGA to calculate the coxph. AML subtype associations are based on z-score enrichment.

OncoSplice further associated each of the novel splicing subtypes with possible splicing regulators by RBP-Finder (**ED Table 2**) or RBP-knockdown correlation. Similar to splice-ICGS, we tested the predictive power of RBP-Finder to identify regulatory RBPs. These predictions include recently experimentally validated regulators, such as MNBL1 as a central regulator of splicing in MLL leukemias^31^. We were able to further obtain strong independent evidence for these predicted regulatory RBPs in 65% of these signatures based on patient genetics (5 out 6) or existing RBP knockdown profiles (8 out 14) (**Supplementary Methods**). Among the novel splicing archetypes detected, nine were also identified in the TCGA AML cohort and six in the TARGET pediatric AML cohort, from independent splice-ICGS analyses (**ED Fig. 2a,b**). We identified dozens of additional known and novel splice architypes in a second large adult AML cohort, BEAT AML, confirming broad and discrete splicing impacts (**ED Fig. 2c**) ^32^.

Strikingly, the two most prominent novel splicing subtypes describe 79% of all AML patients. These subtypes correspond to a single broad splicing-signature, observed when we perform naïve unsupervised clustering of splicing events (**Fig. 1d** and **ED Fig. 1e** ). We herein refer to these two opposing subtypes as *U2AF1-like* and *SRSF2*-like, based on their OncoSplice RBP regulatory predictions and the overlap of these patients with *U2AF1*-S34 and *SRSF2*-P95/8AA mutations, respectively (**Table 1**, **ED Table 2**). Strikingly, this splicing signature was associated with almost 2/3 of all detected splice-events in the entire dataset (∼88,000 events, <u>66%</u>) and could not be described by any known technical effects (e.g., batch, sequence depth), sex or patient age (**ED Table 1**). Notably, however, the incidence of *U2AF1*-like cases was increased (17%) in bone marrow versus peripheral blood, suggesting it may be more indicative of a leukemic stem cell (LSC) signature (**ED Table 1**). When all identified splicing subtypes are directly compared (**Fig. 2b**), our predicted subtypes overlap considerably with several previously established subtypes, suggesting this signature is dominant and can confound the detection of unique splicing in distinct subtypes (**Supplemental Methods – Section 1**). Suprisingly, *U2AF1*-like and *SRSF2*-like splicing account for the large majority of mis-splicing in AML (**Fig. 2c**). We were able to readily confirm these splicing events using read-level visualization (SashimiPlot), including several examples that have been previously reported in unrelated cancer contexts (**Fig. 2d**). A comparative analysis of the frequency of splice-event types from OncoSplice (e.g., cassette-exon versus intron retention) associated with all observed AML subtypes suggests that *U2AF1*-like and *SRSF2*-like have a preference for cassette-exon splicing, similar to that of the majority of RBP mutations, with predictions for other RBPs matching prior literature (e.g., *SF3B1*, *ZRSR2*) ^33,34^ (**Fig. 2e**). To understand the global protein- and domain-level impacts of these splicing event signatures, MultiPath-PSI applies a previously described protein compositional prediction method in the software AltAnalyze ^35,36^. This prediction workflow suggests a significant difference in the potential outcomes of AS, with *U2AF1*-like events shifting isoforms towards full length products that preserve protein-domain integrity while *SRSF2*-like events shift predominantly towards protein truncation or nonsense mediated decay and protein domain disruption (**Fig. 2f**).

### *U2AF1*-like and *SRSF2*-like signatures resolve splicing-factor-mutation specific events

Malignancy associated mutations in SRSF2 and U2AF1 result in gain of function splicing changes, resulting in the selection of atypical splice sites that should not be observed by the wild-type proteoform ^37,38^. However, visualization of the most enriched *U2AF1*-S34 and *SRSF2*-P95 splicing events across all AML indicate that *U2AF1*-like and *SRSF2*-like partially phenocopy mutant-specific splice profiles, respectively (**Fig. 3a,b**). Indeed, approximately half of all alternative splicing events enriched in *U2AF1*-S34 or *SRSF2*-P95 are shared with the respective *U2AF1*-like and *SRSF2*-like signatures when compared to other AMLs (**Fig. 3c**). As previously described, *U2AF1*-S34 has increased specificity for cytosine/adenosine when mediating alternative cassette exon-inclusion at the -3 position of the 3′ splice-site, and a preference for uracil (UAG versus CAG motif) in alternative cassette exon-exclusion (**Fig. 3d**)^39^. To determine whether *U2AF1*-like splicing events where characteristic of wild-type (reference) or mutant *U2AF1*-S34 binding 3’ splice-site sequence recognizition preferences, we performed an enrichment analysis with the software HOMER for unique *U2AF1*-like splicing events (exclusive of *U2AF1*-S34). This analysis finds that while hundreds of atypical 3′ splice-sites are selected in *U2AF1*-like splice events (UAG motif), these represent a fraction of conventional (CAG) events, unlike *U2AF1*-S34 (**Fig. 3d, e**). Thus, while *U2AF1*-like induces a greater number of atypical 3′ splice-sites than *U2AF1*-S34, the pattern of *U2AF1*-like binding remains most similar to wild-type U2AF1, due to the higher number of impacted events (5,193 versus 1,920, respectively). These data support a promiscuous model (over-activation) of U2AF1 directed splicing rather than altered splicing factor specificity in *U2AF1*-like patients.

**Figure 3.**
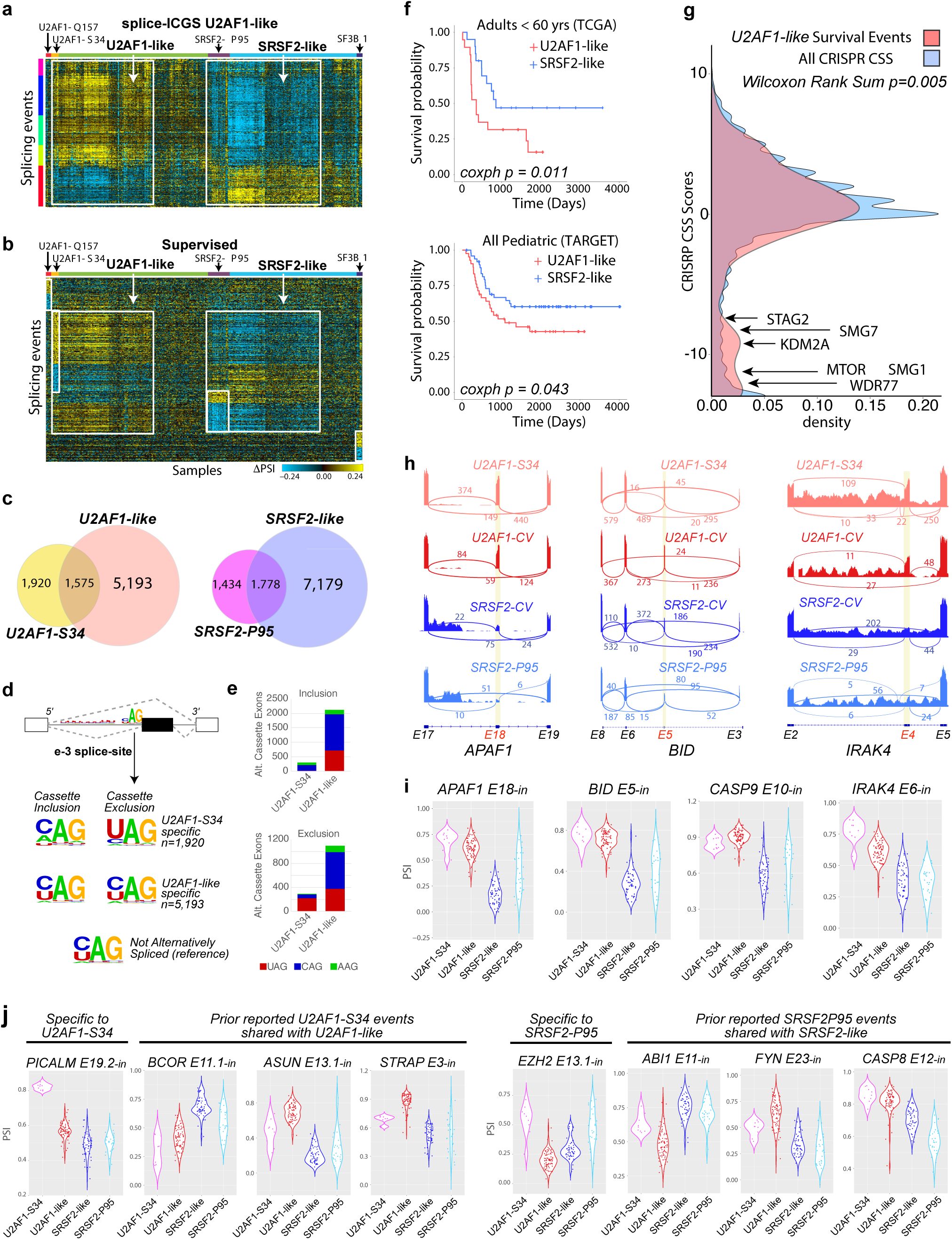
*U2AF1*-like splicing partially phenocopies mutation engendered splicing dysfunction. **a**) splice-ICGS reveals broadly-deregulated splicing in the majority of AML patients. The white boxes indicate 1) RNA-Seq samples with *U2AF1*-S34 mutations and *U2AF1*-like splicing and 2) samples with *SRSF2*-P95 mutations and *SRSF2*-like splicing. **b**). Heatmap showing splicing events enriched (p-value <0.05, FDR adjusted and δPSI =0.1) in adult AML with splicing factor mutations (*U2AF1*-S34, *SRSF2*-P95, *SF3B1*, *U2AF1*-Q157). This supervised analysis identifies the coincidence of *U2AF1*-S34 and *SRSF2*-P95 splicing events with *U2AF1*-like and *SRSF2*-like, respectively (white boxes). **c**) Venn diagram displaying AML-subtype-associated splicing events (MultiPath-PSI) reveals the partial overlap between broadly deregulated and mutation-associated splicing patterns (*U2AF1*-S34 and *U2AF1*-like; *SRSF2*-P95 and *SRSF2*-like). d) Weblogo analysis of U2AF1 binding-site preferences at the e-3 splice-site position for cassette-exon splicing events. *U2AF1*-S34-specific spliced cassette-exons are those not also significant in *U2AF1*-like, while *U2AF1*-like cassette-exons are the inverse. **e**) The number of cassette exon events included and excluded for all *U2AF1*-S34 and all *U2AF1*-like events are shown for each binding site preference. **f**) Kaplan-Meier curves for overall survival in patients from TCGA AML (top) and TARGET AML (bottom) with associated coxph p-values (left: all splice-ICGS stringently classified *U2AF1*-like versus all other considered AMLs. Analysis of TCGA was restricted to cytogenetically normal AMLs with no RNA binding protein (RBP) mutations and under 60 years of age. **g**) Distribution of AML cell-line aggregate CRISPR-screen scores (CSS) of 287 genes corresponding to *U2AF1*-like splicing events (common pediatric and adult) that are significantly associated with poor overall survival compared to all CSS genes. A Wilcoxon rank sum p-value (two-sided) was computed for the comparison of CSS between these two groups. h,i) Select poor-survival associated splicing events in *U2AF1*-like patients and mutation-associated splicing. Example sample SashimiPlots (h) and violin plots of PSI values for all patients (i). j) Violin plot displaying the PSI distribution for previously identified *U2AF1*-S34 or *SRSF2*-P95 splicing events in *U2AF1*- and *SRSF2*-like patients. in=inclusion exon, ex=exclusion exon.

### *U2AF1*-like splicing is associated with poor survival and leukemic growth

Although no *U2AF1* or *SRSF2* mutants were observed in pediatric AML, we find the same splicing signature with splice-ICGS in TARGET AML patients (**ED Fig. 3a**). Strikingly, in both TCGA and TARGET, *U2AF1*-like splicing events are associated with poor overall survival (cox proportional hazard (coxph) p = 0.01 and 0.04, respectively), while *SRSF2-like* splicing events are associated with improved survival in TCGA but not in TARGET (coxph p-value = 0.02 and 0.08, respectfully) (**Fig. 3f**). *U2AF1*-like was further associated with decreased time to relapse in adult but not pediatric AML (coxph p=0.02). To determine whether *U2AF1*-like and *SRSF2*-like biology can be modelled *in vitro*, we analyzed bulk RNA-Seq on pediatric AML patient blasts at the time of diagnosis and upon cell culture (**ED Fig. 3b**) ^40^. While diagnostic cells were a mix of *U2AF1*-like, *SRSF2*-like and intermediate cells based on RNA-Seq, all cells upon culturing transitioned to a *U2AF1*-like profile. Further projecting these splicing signatures across all leukemic cell lines with RNA-seq in the CCLE project^41^ also finds strongly skewed splicing towards the *U2AF1*-like (**ED Fig. 3c**). Hence, *U2AF1*-like (but not *SRSF2*-like) liabilities can be assessed in immortalized cells, with the splicing choice in these cells likely to be mediated by extrinsic factors (e.g., stromal niche, cytokines).

Given that *U2AF1*-like impacts can be assessed in vitro, to understand the functional consequences of genes mis-spliced in *U2AF1*-like patients, we restricted events to those *U2AF1*-like splicing events both associated with poor overall survival (n=287 genes, TARGET coxph p<0.05) and shared in pediatric and adult patients and assessed these in a prior broad CRISPR dependency screen in 13 CRISPR AML cell lines ^42^. Among these 287 genes, 42 were required for leukemic growth in >6 cell lines, greater than expected by chance (Wilcoxon rank test p<0.005, two-sided) (**Fig. 3g**). Thus, *U2AF1*-like splicing events correlated with patient survival are enriched for genes required for leukemic growth.

To understand the broader significance of these observed splicing events, we compared *U2AF1*-like splicing events to prior-described cancer-associated splice isoforms and genes likely to impact key cancer pathways. Notably, multiple well-characterized cancer associated splicing events were found that have a graded response in the mutants and “like” patient subsets (*APAF1*, *BID*, *CASP9, FAS*) in addition to well-defined therapeutic cancer targets (*MTOR*, *KDM2A*, *MAPK14*, *IL6R*, *TLE4*, *IRAK4*) (**Fig. 3h,i** and **ED Fig. 3d-f**). This data further agree with our prior observation that the highest inclusion of exon 4 IRAK4 is coincident with the U2AF1-S34 mutation, but with many patients exhibiting a similar graded pattern in exon 4 splicing. Many of the identified *U2AF1*-like regulated splicing events also include prior annotated targets of mutant *U2AF1* and *SRSF2* in AML and MDS, with important well-documented exceptions for *SRSF2*-P95 (e.g., *EZH2*) and *U2AF1*-S34 (e.g., *PICALM*) targets ^37–39,43,44^, which are not found within the “like” signatures (**Fig. 3j**). Thus, consideration of *U2AF1*-like splicing informs the specificity of mutant-specific events.

### *U2AF1*-like is stable overtime and derives from a circadian splicing program

To confirm that *U2AF1*-like splicing is durable and contributes to prognosis over time, we re-analyzed an independent set of serial diagnosis and relapse AML RNA-Seq of 19 patients ^45^. Patients with annotated *U2AF1*-like or *SRSF2*-like AML showed the same pattern at diagnosis and relapse when considering (**Fig. 4a**). *U2AF1*-like and *SRSF2*-like patterns were independent from prior-defined cytosine-methylation profiling-defined subtypes in this cohort, suggesting that mis-splicing is not a biproduct of broader epigenetic alternations ^45^. We confirmed the overall stability of *U2AF1-like* splicing from 41 patients with 2-3 samples at independent timepoints of therapy or relapse in BEAT-AML (**Fig. 4b**).

**Figure 4.**
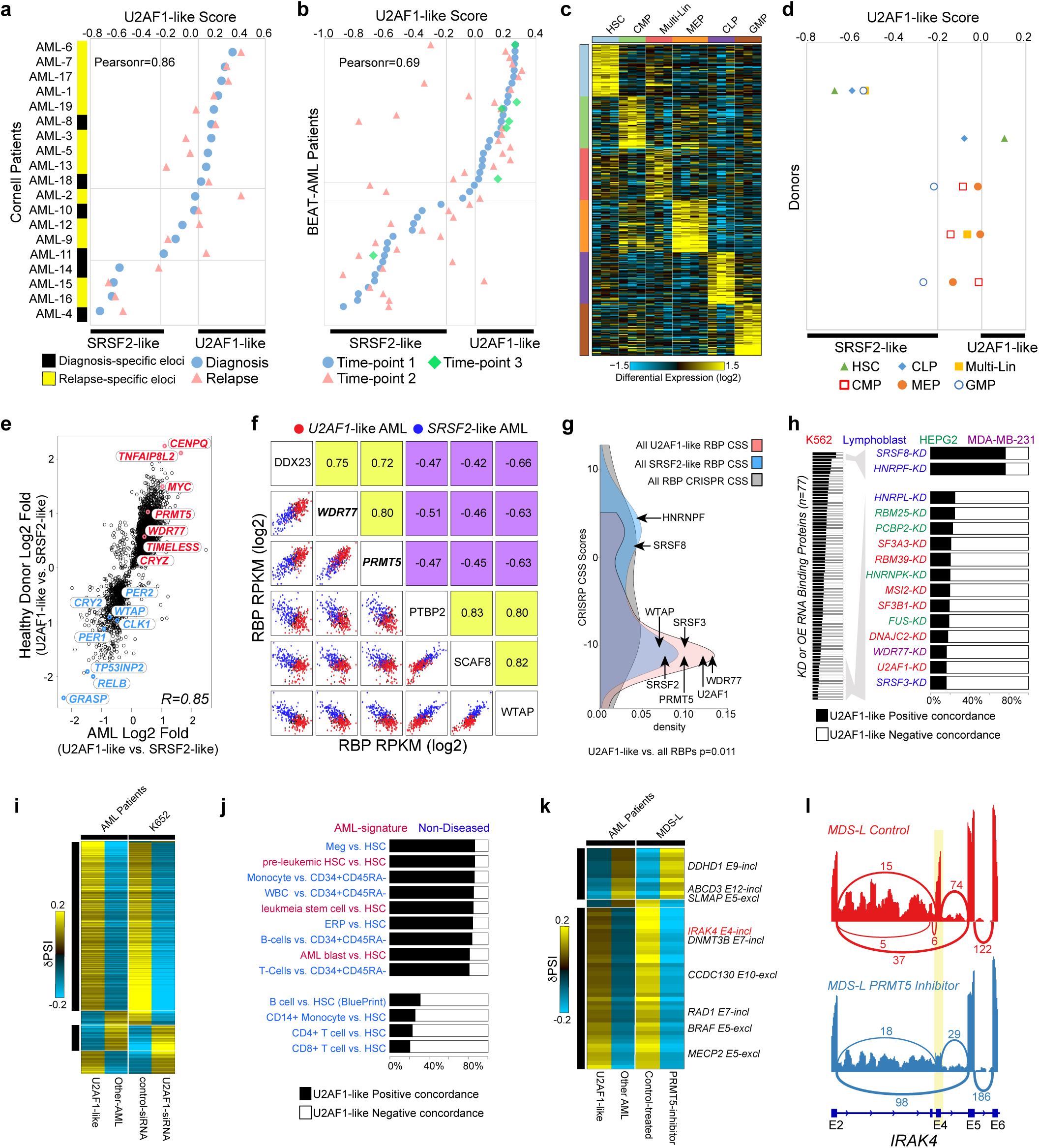
*U2AF1*-like splicing is mediated by PRMT5 and WDR77 expression. **a**) *U2AF1*-like splicing status in a prior relapse cohort of matched AML samples at diagnosis (blue dot) and relapse (pink triangle)^45^. The *U2AF1*-score is calculated from the aggregate of 364 poor survival-associated *U2AF1*-like inclusion versus exclusion ΔPSI splicing event values for each patient. Classification of AML samples as *U2AF1*-like, *SRSF2*-like, or Other are based on the range of scores produced for Leucegene patients assigned by splice-ICGS. Prior annotated epigenetic subtypes (eloci) are indicated (below). **b**) *U2AF1*-like splicing status in AML patients from with multi-timepoint sampling in the BEAT-AML RNA-Seq cohort. **c**) Heatmap of the top marker genes (MarkerFinder) distinct human hematopeoitc stem and progenitor populations isolated by Fluorescence-Activated Cell Sorting (FACS) (BluePrint consortium). **d**) *U2AF1*-like scores in the BluePrint RNA-Seq segregate by donor rather than cell-type. **e**) Scatter plot comparing gene expression for 3,574 commonly differentially expressed genes between *U2AF1*-like and *SRSF2*-like samples matched in the AML and healthy donor datasets (eBayes t-test p<0.05 (two-sided)). Select transcription factor, splicing regulator and circadian regulators are denoted. **f**) Top-associated *U2AF1*-like splicing factors, by comparing gene expression and splicing (Pearson correlation). Correlation values are shown in the upper right quadrants. Scatter plots (lower left quadrants) illustrate the pairwise expression value of the indicated factors. **g**) AML cell-line CRISPR-screen scores (CSS) for RBPs associated with *U2AF1*-like or *SRSF2*-like splicing from gene expression or knockdown signature analyses compared to all RBPs. **h**) The extent of splicing concordance (similarity index) between Leucgene AML *U2AF1*-like splice events to RBP knockdown (KD) or over-expression (OE) (n=77) in the indicated cell lines. i) Heatmap of all AML *U2AF1*-like significant splicing events overlaping to shRNA KD of *U2AF1* in K562 cells (ENCODE). **j**) Concordance between AML *U2AF1*-like splice events with cell-type and hematological malignancy specific splicing programs. **k**) Heatmap of all AML *U2AF1*-like significant splicing events overlapping with PRMT5 inhibitor treated MDS-L cell RNA-Seq. **l**) SashimiPlot of the IRAK4 gene locus in PRMT5 inhibitor treated and control MDS-L cell RNA-Seq (prior annotated Exon 4 (denoted E6 in AltAnalyze)).

To determine if *U2AF1*-like splicing is simply indicative of distinct progenitor populations, we evaluated RNA-Seq from different sorted cord blood CD34+ cells (HSC, MPP, CMP, GMP, CLP, MEP, Multi-Lin)^46^. Surprisingly, these analyses find that *U2AF1*-like splicing differs by donor rather than by progenitor cell-type (**Fig. 4c,d** and **ED Fig. 4a**). Indeed, comparison of *U2AF1*-like splicing in healthy donor bone marrow CD34+ progenitors finds a separation of *U2AF1*-like and *SRSF2*-like splicing patterns in healthy donors, although for a fewer subset (1/6^th^) of the AML splicing events (1139/6769) (**ED Fig. 4b,c**) ^47^. While only a small proportion of *U2AF1*-like splicing events from AML are present in healthy donors, concordance in the gene expression profiles of AML and healthy CD34+ cells were high between AML and healthy bone morrow associated *U2AF1*-like architypes (97% agreement). Comparing AML and healthy progenitors, *MYC* was among the most consistent induced gene in *U2AF1*-versus *SRSF2*-like patient/donors, along with multiple splicing regulators (e.g., *WDR77*, *PRMT5*, *WTAP, CLK1*) (**Fig. 4e**). These shared differentially expressed genes were notably enriched in core regulators of circadian rhythm (e.g., *ARNTL, PER1, PER2, CRY2, NCOR1, PRMT5*) (**ED Table 3**). To determine whether *U2AF1*-like splicing in AML mimics a normal circadian splicing signature, we applied the machine learning program CYCLOPS (cyclic ordering by periodic structure) ^48^ to the AML RNA-Seq gene expression data. When visualizing both circadian predicted phase ordering and splicing-defined patient groups, we find *U2AF1-like* and *SRSF2*-like samples can be predicted from circadian phase alone **(ED Fig. 4e**). These two major circadian orded phases were primarily associated with extracellular signaling/inflammatory (*SRSF2*-like) versus proliferative/metabolic (*U2AF1*-like). Thus, *U2AF1*-like is associated with physiological circadian splicing, which is predictive of patient overall survival.

### *U2AF1*-like splicing is regulated by a MYC driven WDR77/PRMT5 splicing program

While the association with circadian rhythm is intriguing, it does not explain what the drivers of mis-splicing are in AML. We find *MYC* to be among the most significantly upregulated genes in *U2AF1*-like patients (p=1e-13) along with several previously validated MYC ^49^ and PRMT5/WDR77 ^50,51^ dependent splicing events, based on re-analysis of these published data (**ED Fig. 4f,g**). MYC has been shown to regulate PRMT5 and WDR77 expression as well as the splicing of several pro-oncogenic splicing events ^51^. In both Leucegene and TARGET, OncoSplice consistently identified correlated or anti-correlated expression and significant differential expression of multiple splicing factors with *U2AF1*-like splicing, notably the cancer splicing modulators *WDR77*, *PRMT5* and *WTAP* ^10,49,51^ (**Fig. 4f**). WDR77 and PRMT5 form a coactivator complex that dimethylates specific arginines in several spliceosomal Sm proteins, resulting in alternative splicing ^50^. In the AML CRISPR dependency data, the *U2AF1*-like associated RBPs, *U2AF1*, *SRSF2*, *WDR77* and *PRMT5,* were predicted to be essential in AML compared to all targeted genes (p=0.011) (**Fig. 4g** and **ED. Fig. 4i**). To independently assess splicing factor regulatory potential for *U2AF1*-like splicing, we analyzed a large repository of RBP KDs from ENCODE and prior and diverse published studies. Consistent to our differential expression and CLIP-Seq analyses, *U2AF1* and *WDR77* knockdown effectively reversed *U2AF1*-like splicing (>80% negative concordance), along with *SRSF3* (**Fig. 4h,i** and **ED Fig. 4h**). Knockdown of *SRSF8* and *HNRNPF* had the opposite impact, resulting in a shift towards *U2AF1*-like spliced isoforms. Expression of *WDR77* alone was found to be a novel independent predictor of patient survival when considering both TCGA and TARGET (**ED. Fig. 4j**).

Our gene expression and CRISPR screen comparative analyses, suggest *U2AF1-like* and associated upstream RBPs mediate a proliferative and stem cell maintenance program, while *SRSF2-like* lacks this program, but is associated with inflammation. To initially understand this relationship, we extended our splicing signature comparisons to previously described cell-type specific and LSC splicing comparison RNA-Seq datasets ^52^. This analysis finds that *U2AF1-like* strongly phenocopies splicing in LSC versus HSC (**Fig. 4j**). This finding was unique to LSC comparisons, versus alternative hematopoietic cell-type comparisons, suggesting that *U2AF1*-like drives a core stem cell program which is lacking in *SRSF2*-like.

PRMT5 inhibition using selective small molecule inhibitors have emerged as a promising strategy to inhibit leukemic cell growth in some hematological malignancies ^53^. To determine whether PRMT5 inhibition specifically impacts *U2AF1*-like as opposed to *SRSF2*-like splicing, we next performed RNA-Seq in the AML cell line, MDSL ^54^, treated with a specific PRMT5 inhibitor. While the PRMT5 inhibitor primarily induced intron retention (474/720 unique PSI events), PRMT5 blockade reversed the *U2AF1*-like splicing for 90% of exonic splicing events (**Fig. 4k**). The previously identified *IRAK4* exon 4 inclusion (associated with hypermorphic IRAK4-Long) was among the top rescued *U2AF1*-like splicing events (**Fig 4l**). These findings implicate PRMT5 as a regulator of IRAK4 isoforms in MDS/AML.

### PRMT5 inhibition suppresses IRAK4-L expression and leads to increased myeloid differentiation and impaired MDS/AML progenitor cell function

Based of these data, we hypothesized that PRMT5 regulation of the IRAK4-Long(L) isoform promotes a LSC maintenance program in *U2AF1*-like MDS/AML that blocks differentiation. In MDS-L cells, PRMT5 inhibition led to reduced expression of IRAK4-L as seen by immunoblotting, suggesting that the exclusion of IRAK4 exon 4 by PRMT5 results in reduced IRAK4-L protein expression (**Fig 5a**). Consistent with this reduction in IRAK4-L protein, PRMT5 inhibition led to decreased activity of NF-kB in a myeloid leukemia cell line reporter assay (**Fig. 5b**). Suppression of IRAK4-L and NF-kB coincided with a significant dose-dependent decrease in viability of PRMT5 inhibitor-treated MDSL cells (**Fig. 5c**). The loss of viability was pronounced with longer duration of treatment and was accompanied by myeloid differentiation as evident from cytomorphology and increased expression of CD14 and CD11b in FACS analysis (**Fig. 5d,e**).

**Figure 5:**
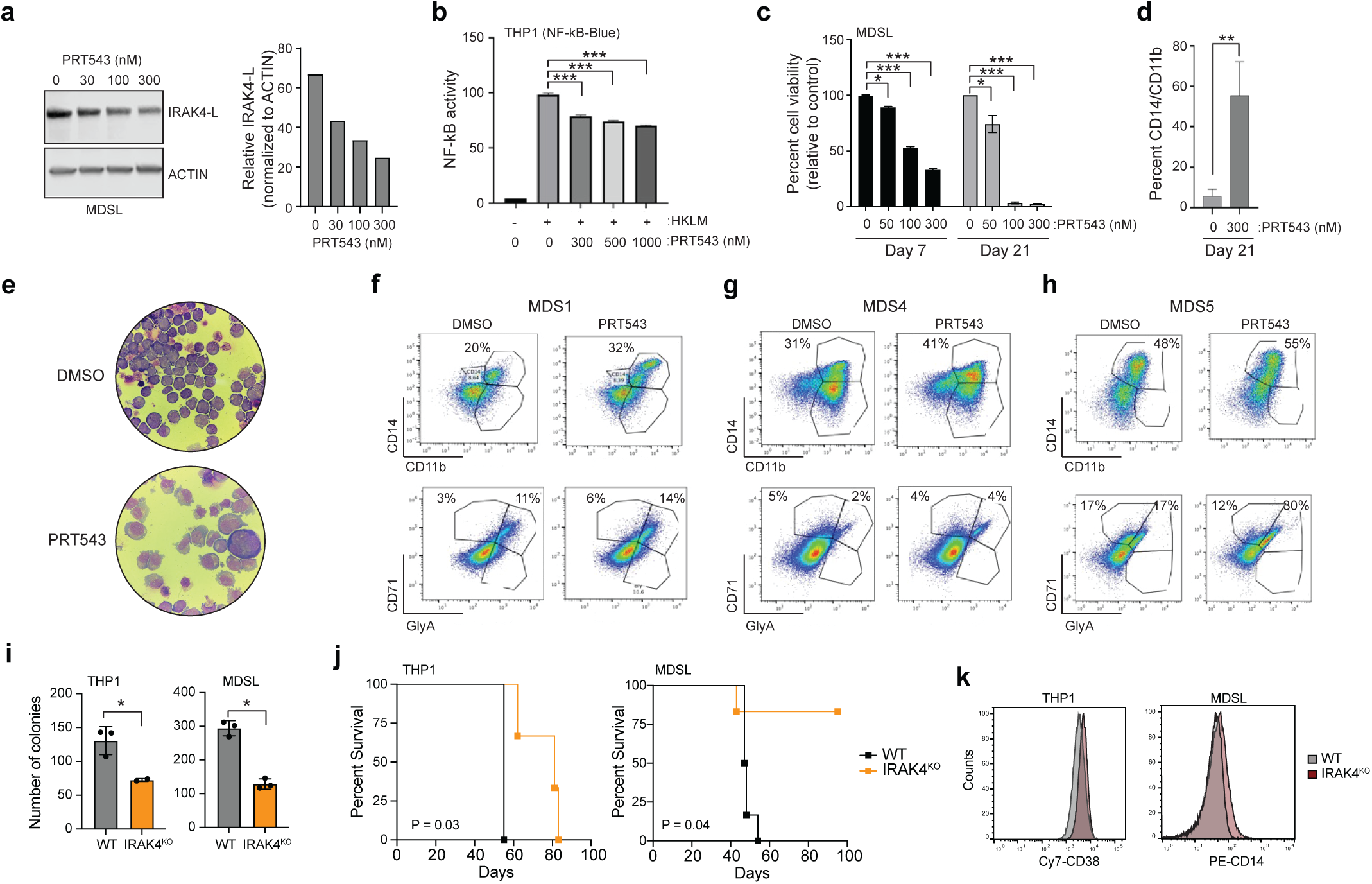
Treatment with PRMT5 inhibitor PRT543 via targeting of IRAK4 leads to increased myeloid differentiation and impaired MDS/AML progenitor cell function. **a**) Western blot of IRAK4-L (long isoform) in MDSL cells following 72 hours of treatment with the PRMT5 inhibitor (PRT543) at 30nM, 100nM and 300nM. **b**) NF-kB reporter activity in THP1-Blue NF-kB cells with increasing doses of the PRMT5 inhibitor (PRT543). **c**) Cell viability in MDSL cells with increasing concentrations of PRT543 50nM, 100nM and 300nM as compared to control. **d**) MDSL were treated with PRT543 300nM every 72 hours as compared to control and myeloid differentiation assessed on days 21. Statistically significant differentiation occurred in the treated cell population as compared to control at day 21 for CD14 + CD11b. (N=3, P <0.01). **e**) Representative images of Giemsa stained MDSL cells alone and those treated with PRMT5 inhibitor (PRT543) at 300nM treatment for 72 hours. The red arrow identifies evidence of differentiated myeloid cells with red arrowhead showing evidence of increased vacuolization, consistent with myeloid maturation/differentiation. **f-h**) MDS patient samples treated with PRMT5i and control for 14 days in clonogenic assays and then assessed for myeloid and erythroid differentiation by FACS. **i**) Colony formation in isogenic WT and IRAK4^KO^ THP1 and MDSL cell lines (two independent experiments). **j**) Kaplan Meier survival analysis of NSGS mice (n = 5 mice/group) engrafted with WT or IRAK4^KO^ THP1 or MDSL cells (Data represent one of two independent experiments with similar trends). **k**) Immunophenotyping of the indicated cells for CD38 and CD14 expression, respectively.

Myeloid differentiation blocks are a hallmark of MDS/AML and IRAK4-L has been shown to be expressed preferentially in MDS patient blasts ^13^. To understand the effects of PRMT5 inhibition in MDS samples we cultured primary patient bone marrow samples with the PRMT5 inhibitor (PRT543) to assess for functional activity. Supporting our hypothesis, treatment with PRT543 led to increased myeloid differentiation of primary MDS samples (**Fig. 5f-h**). These data demonstrate the preclinical efficacy of targeting PRMT5 in myeloid malignancies. Recent studies have implicated the IRAK paralogs, IRAK1 and IRAK4, in MDS/AML by preserving the undifferentiated state of LSCs ^55^. To confirm whether the expression of IRAK4-L is critical to the maintenance of LSC fitness, we evaluated a panel of isogenic AML cell lines in which the IRAK4 was deleted using CRISPR/Cas9 editing. In agreement with induced expression of IRAK4-L in *U2AF1*-like AML, deletion of IRAK4 in MDS/AML cells (IRAK4^KO^) resulted in reduced colony formation *in vitro* (**Fig. 5i**) and leukemia development in xenografted mice (**Fig. 5j**). To determine whether IRAK4 is required for preserving an undifferentiated LSC state, we examined morphological changes of MDS/AML cells upon deletion of IRAK4. In contrast to WT cells, IRAK4^KO^ AML cells exhibited increased expression of differentiation markers (**Fig. 5k**), which is congruent with myeloid differentiation. These findings suggest that PRMT5 mediates IRAK4-L expression and that IRAK4 function is important for preserving an immature cell state of LSCs.

## DISCUSSION

While the genetic evaluation of cancers has significantly improved cancer risk stratification and treatment, patients with a similar constellation of mutations frequently have widely varying outcomes. Unsupervised evaluation of molecular subtypes in diverse diseases, particularly in cancer, has revealed novel disease states and therapeutic targets ^56,57^. Using an iterative unsupervised approach for splicing-archetype discovery (splice-ICGS), we find clinically significant archetypes that occur independently of observed tumor genetics and predominant tumor gene expression patterns. In addition to finding over a dozen genetically-defined subtypes of AML, OncoSplice identified two major AML populations defined selectively by alternative splicing, *U2AF1*-like and *SRSF2*-like, which together describe close to 80% of all adult and pediatric AMLs. Our analyses further suggest that non-pathogenic *U2AF1*-like splicing in healthy bone marrow, which comprises only 16% of the events found in AML, predisposes hyper-active U2AF1 signaling upon leukemic cell transformation and poor survival. Finally, we show that the core splicing program of *U2AF1*-like patients can be rescued pharmacological inhibition of its implicated regulator PRMT5 and implicate its target, IRAK4, as mediator LSC maintenance.

The identification of novel splicing subtypes has additional important implications for the understanding of complex diseases, such as cancer. In this study, *U2AF1*-like and *SRSF2-like* splicing clarifies highly specific splicing events associated with mutant versions of those proteins. Notably, alternative splicing of *IRAK4* was recently proven to be a therapeutic vulnerability in MDS and AML; however, while *U2AF1*-S34 expression induced *IRAK4* exon-inclusion, AML without *U2AF1* mutations also showed the therapy-relevant splice event ^13^. In both scenarios, IRAK4 exon-inclusion resulted in the expression of a hypermorphic isoform that is capable of signaling in the absence of upstream receptor activation ^14^. We find that the same long isoform of *IRAK4* is among the most enriched *U2AF1*-like enriched splicing-events, repositioning *IRAK4* as a therapeutic target for the *U2AF1*-like-defined AML patient population. Importantly, the IRAK4-Long isoform is expressed at lower levels in normal HSCs ^13^. Given the emerging role of IRAK4 signaling in human diseases, IRAK4 inhibitors and proteolysis targeting chimeric (PROTAC) small molecule degraders are being assessed in pre-clinical studies and clinical trials for hematologic malignancies and inflammatory conditions. Additionally, using splicing as a readout enables the separation of patients with a similar spectrum of mutations (*NMP1, TP53, FLT3-ITD*) or the unification of patients with diverse mutations within the same genes that have a common splicing profile (*SRSF2, HNRPNK, ZRSR2, SF3B1*). Differential splicing observed in both adult and pediatric AMLs was highly concordant, suggesting common therapeutic vulnerabilities. The identification of these new subtypes provides opportunities for identifying patients likely resistant to therapy and proposes selective strategies for emerging therapeutic targeting (i.e., PRMT5/WDR77 or IRAK4) ^58^. Application of these unsupervised computational approaches beyond leukemia and even beyond splicing (**ED Fig 3c-d**) is likely to shed new light on tumor heterogeneity and the pathways that underlie therapeutic response.

## METHODS

### RNA-Sequencing

RNA-Seq of non-diseased bone marrow was performed from donors who have been consented under the IRB-approved Normal Donor Repository (Cincinnati Children’s Hospital Medical Center). RNA was processed from young adult healthy CD34+ bone marrow. The total RNA was extracted by using mirVana miRNA Isolation Kit (Lifetech, Grand Island, NY) with total RNA extraction protocol. In brief, freshly prepared cells were immediately lyzed by Lysis/Binding Buffer, treated with Homogenate Additive, and followed by Acid-Phenol:Chloroform extraction according to the standard protocol. The supernatant was mixed with ethanol and passed through Filter Cartridge. The bound RNA was then washed and eluted. The RNA concentration was measured by Nanodrop (Thermo Scientific, Wilmington, DE) and its integrity was determined by Bioanalyzer (Agilent, Santa Clara, CA). Sequencing was performed on an Illumina HiSeq 1000 using single-end sequencing at a target depth of 20 million reads per sample. The RNA-Seq data were processed using the same alignment workflow applied to the primary AML samples. These data have been deposited in the Gene Expression Omnibus (GSE118944).

### RNA-Seq Quantification and Variant Analysis

Primary adult AML RNA-Seq FASTQ files were obtained from the Leucegene consortium (GSE67040, GSE62190, GSE49642) and re-processed using STAR to hg19, allowing for the identification of known (UCSC mRNAs) and *de novo* junctions for the same samples. STAR was used in concert to identify predicted sequence deletions (SRSF2-8AA del). TCGA tier-1 and BEAT-AML adult AML and TARGET pediatric AML RNA-Seq samples were obtained from the Genome Data Commons following controlled access approval from dbGaP and processed using the same alignment options. Normal donor bone marrow progenitor RNA-Seq was obtained from GSE63569 ^59^. ENCODE knockdown raw RNA-Seq FASTQ files were obtained from the ENCODE project (https://www.encodeproject.org/matrix/?type=Experiment&status=released&assay_title=shRNA+RNA-seq&assembly=hg19&target.investigated_as=RNA+binding+protein&biosample_ontology.organ_slims=blood). Gene expression was quantified with AltAnalyze version 2.1.1 default RPKM analysis pipeline. Spliced exon-exon and exon-intron junction reads were quantified in AltAnalyze using the MultiPath-PSI method in conjunction with AltAnalyze’s BAM file intron quantification module (BAMtoExonBED). MultiPath-PSI examines each known and novel exon-exon or known exon-intron junction in a sample and computes its relative detection compared to the local background of all genomic overlapping junctions that can be directly associated with the given gene. This algorithm employs the same statistical approach to identify high confidence intron retention events but evidenced by pairs of exon-intron and intron-only mapping paired-end reads, sufficiently detected at both ends of a given intron (5′ and 3′). Additional details can be found in the Supplemental Methods. Splicing-event annotation types (e.g., cassette-exon, alternative 5’ splice-site, intron retention) and domain-level functional consequences were automatically supplied by AltAnalyze. Conventional gene-set enrichment analyses were performed using the GO-Elite algorithm in AltAnalyze ^60^.

### OncoSplice algorithm

Full details regarding methods implemented, validation and background information in the full OncoSplice pipeline can be found in Supplemental Methods.

### Identification of Cancer Genomic Variants

For the Leucegene dataset, genome variants were detected using the GATK RNA-Seq analysis workflow ^61^ and annotated through STAR insertions/deletions ^62^, COSMIC ^63^ and Ensembl Variant Effect Predictor ^64^ (**Fig. S1b**). Oncofusions were detected with the rigorous FusionCatcher pipeline ^65^. Additional variant and clinical annotations were obtained from the TCGA and TARGET consortiums and where available from previously described Leucegene subsets or from the MISTIQ database where available (initially blinded from our analysis) ^22,66,67^. Variant and oncofusion enrichment analyses were performed using a Chi-squared test (p<0.05) aggregating variants at the gene level. Disease free and overall survival analyses were performed in R using the multivariate cox proportional hazard (coxph) tests for each splicing subtype. The R packages glmnet and coxph were used to test for other clinical covariates such as subtype/grade, cytogenetic abnormalities, relapse, induction failure or secondary site of metastasis, while accounting for potential confounding variables such as age, gender, ethnicity, smoking, drug therapy or subtype/grade. Co-occurring genomic variant or common splicing events between subtypes were visualized using Circos plots with the circos package ^68^. Using GATK pipeline, KM analyses and literature data were integrated to identify patients with common mutations in AML. Enrichment analyses were assessed using Fisher’s Exact Test p-values following FDR correction. In addition, z-scores, sensitivity and specificity were also calculated for each mutation and the different splicing subtypes to find any associations. Further, these analyses were extended to identifying co-occurring mutation enrichments.

### Human Patient Samples, Cell lines and Reagents

Patients diagnosed with MDS were obtained after IRB approval by the Albert Einstein College of Medicine. The AML cell line MDS-L was provided by Dr Starczynowski ^54^ and was cultured with the addition of 10ng/ml human recombinant IL-3. THP1 were purchased from the American Type Culture Collection. THP1 were cultured in RPMI-1640 medium with

10% FBS and 1% penicillin–streptomycin. The THP1 IRAK4^KO^ and MDSL IRAK4^KO^ clones were previously described ^55^. PRMT5 inhibitor PRT543 was obtained from Prelude Therapeutics.

### NF-kB Reporter Assays

The reporter cell line-THP1-Blue (TM) NF-kB SEAP reporter (Invivogen, Cat# thp-nfkb) derived from the human THP1 monocytes cell line was obtained. This assay used Heat Killed Listeria Monocytogenes (HKLM), a TLR2 agonist that triggers the NF-kB pathway. THP1-Blue NF-kB cells were seeded in a 96 well plate at a density of 2 x 10^4^ cells /well for the following conditions 1) Cells alone control 2) HKLM alone control 3) PRT543 - 50nM 4) PRT543 - 300nM and 5) PRT543 - 1000nM. Following 24 hour incubations, cell supernatants were assayed with Quanti-Blue medium for 4-6 hours according to the manufacturer’s instructions and the levels of NF-kB induced SEAP were detected at 650 nM using the Fluostar Omega Microplate reader.

### Cell Viability assays

Cell viability assay was performed using Cell titer blue (Promega, Madison, WI). MDS-L cells were seeded in a 96 well plate at a density of 5000 cells / well and treated with different concentrations of PRT543 ranging from 50nM, 100nM to 300nM. Dosing was done starting from Day 0 and continued on every third day. Day 7 plate receiving a total of 3 doses and Day 21 plate a total of 7 doses, were assessed for cell viability by the addition of cell titer blue (Promega). Fluorescence was measured using the Fluostar Omega Microplate reader (BMG lab tech).

### Flow cytometry analysis for Myeloid differentiation markers

MDS-L cells were seeded in a 6 well plate at a density of 100,000 cells and treated with PRT543 at a concentration of 300nM. Dosing was done starting at Day 0 and continued every third day (total of 7 doses). On Day 21, cells were stained with Human CD11b-APC conjugate ( Thermo Fisher Scientific, Waltham, MA, USA, Catalogue No CD11B05, clone VIM12), Human CD14-Pacific blue TM ( Thermo Fisher Scientific, Catalogue No MHCD1428, clone Tuk 4). Using a BD FACS LSRII instrument ( BD Biosciences, Franklin Lakes, NJ, USA) data was acquired and analyzed using Flow Jo software version 10.6.1 (BD Biosciences).

### Clonogenic Progenitor assays

Primary patient MDS samples were plated in Methylcellulose ( Stem cell technologies, H4435, Vancouver, CA) with PRT543 at different concentrations and control and colonies were counted after 14 -17 days. This was followed by staining and processing by Flow cytometry (BD FACS LSRII instrument) for Erythroid and Myeloid differentiation. Antibodies used were Human CD45 PE-Cy7; Human CD34 PE; Human Gly-A PerCP-Cy5.5; Human CD14 Pacific blue; Human CD71 FITC and Human CD11b APC. For THP1 and MDSL clonogenic assays, clonogenic frequencies were determined by plating cell lines in Methocult H4434 (StemCell Technologies) in SmartDish meniscus-free 6-well plates (StemCell Technologies). Plates were kept in humidified chambers and colonies were imaged and manually scored after 9-14 days using the STEMvision counter (StemCell Technologies).

### GIEMSA staining

MDS-L cells control and PRT543 treated (3 doses) were cytospun on slides and stained with Giemsa solution.

### Western blot analysis

MDS-L cells control and PRT543 treated were harvested and protein lysates incubated for 30 minutes with western lysis buffer, containing cocktail phosphatase inhibitors and proteases. Immunoblotting was performed by LI-COR western blotting using IRAK-4 antibody to demonstrate reduction of the oncogenic IRAK-4 signaling pathway and confirm a decrease in IRAK-4 Long isoform.

### Xenografts

Animals were bred and housed in the Association for Assessment and Accreditation of Laboratory Animal Care-accredited animal facility of Cincinnati Children’s Hospital Medical Center (IACUC2019-0072). For the xenograft using isogenic THP1 or MDSL cells, WT or IRAK4^KO^ cells were suspended in PBS and injected via tail vein into NOD.Cg-*Prkdc^scid^ Il2rg^tm1Wjl^* Tg(CMV-IL3,CSF2,KITLG)1Eav/MloySzJ (NSGS) mice at a dose of 2.5 x 10^5^ cells per mouse. Moribund mice were sacrificed and assessed for leukemic burden measurements. Briefly, mice were euthanized with carbon dioxide following the AVMA Guidelines for the Euthanasia of Animals and BM cells were immediately extracted by breaking the femurs with a mortar and pestle. BM cells were frozen in FBS with 10% DMSO until the time of analysis. BM was analyzed for huCD45 (BDPharmingen, Cat#555485) and huCD33 (BDPharmingen, Cat#555450) expression by flow cytometry using a BD LSRFortessa (BD Biosciences). For staining, 1x10^6^ cells from each BM sample were incubated with antibodies diluted 1:100 in a solution of PBS, 0.2% FBS for 30 minutes on ice in the dark. Cells were washed once with PBS, resuspended in PBS with 0.2% FBS, and immediately analyzed by flow cytometry.

### Statistical analysis

For non-genomic analyses, differences among multiple groups were assessed by one-way analysis of variance (ANOVA) followed by Tukey’s multiple comparison posttest for all possible combinations. Comparison of two group was performed using the Mann-Whitney test or the Student’s *t* test (unpaired, two tailed) when sample size allowed. Unless otherwise specified, results are depicted as the mean ± standard deviation or standard error of the mean. A normal distribution of data was assessed for data sets >30.

## ACKNOWLEDGMENTS

This work was supported by Cincinnati Children’s Hospital Research Foundation, funding from the Edward P. Evans Foundation’s EvansMDS initiative, the National Institutes of Health R01 CA196658 (HLG), R01 CA226802 (NS), and R01 NS099068 (MW), R01CA275007 (AV and DTS), CCRF Endowed Scholar (MTW), CCHMC CpG Pilot study award (MTW, NS), Prelude Therapeutics, CCHMC Trustee Awards (MTW, NS), R35HL135787 (DTS), and Cancer Free Kids (DTS, HLG). This work was supported by NIDDK U54 DK126108 at Cincinnati Children’s Hospital Medical Center and their Flow Cytometry and Comprehensive Mouse Cores. We thank J Bailey and V Summey for assistance with transplantations (CCHMC Comprehensive Rodent and Radiation Facility; RRID:SCR_022624).

## AUTHOR CONTRIBUTIONS

The manuscript was written by MV, MW, HLG, AV and NS. Experiments were designed by DS, AV, HLG and NS, and performed by CH, AO, TN, LS, NR, SGM, AF, DTS, SS and KM. Reagants supplied by PS and DH. Computational analyses were performed by MV, KC, AK, XN, XC, MW, KP and NS. NS, AV, DTS and HLG interpreted data and wrote and/or edited the manuscript. All authors approved the final version of the manuscript.

## SUPPLEMENTAL FIGURE LEGENDS

**ED Figure 1.**
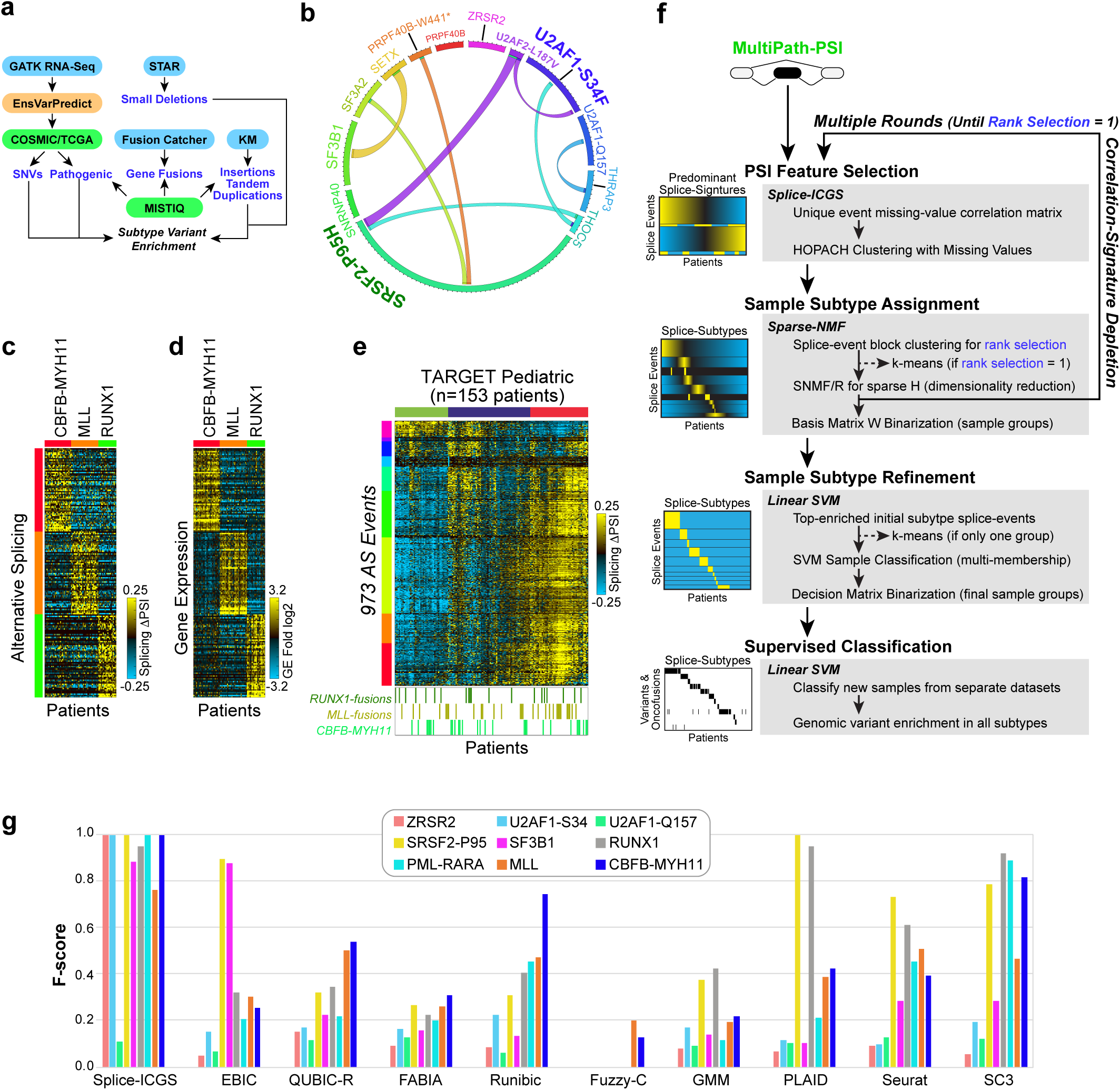
In silico detect and evaluation of splicing-versus gene-expression defined molecular subtypes. **a**) Schematic of the computational workflow to define genomic variants (GATK), oncofusions (Fusion Catcher), insertions and tandem duplications (KM), small gene deletions (STAR) and variant annotation (COSMIC, TCGA, EnsVarPredict, MISTIQ) in the Leucegene cohort. **b**) Circos plot depicting the relative frequency and pairwise co-occurrence of splicing factor mutations implicated in AML and MDS that were detected in RNA-Seq (Leucegene). **c,d**) Heatmap of the top marker alternative splicing events (c) or differentially expressed genes (d) using the MarkerFinder algorithm for common pediatric onocofusions (TARGET AML cohort). **e**) Heatmap of predominant splicing patterns in TARGET AML RNA-Seq using the software ICGS. Colored bars indicate patients with known mutations or oncofusions (below). **f**) Prinicipal algorithm steps in the interative splice-ICGS workflow to identify partially overlapping splicing-defind molecular subtypes. **g**) Performance of splice-ICGS compared to well-described approaches for unsupervised molecular subtype identification for bulk and single-cell transcriptomics data. Performance is measured according to F-score (harmonic mean of precision and recall) for each genetically defined subtype of AML evaluated in Leucegene (PSI).

**ED Figure 2.**
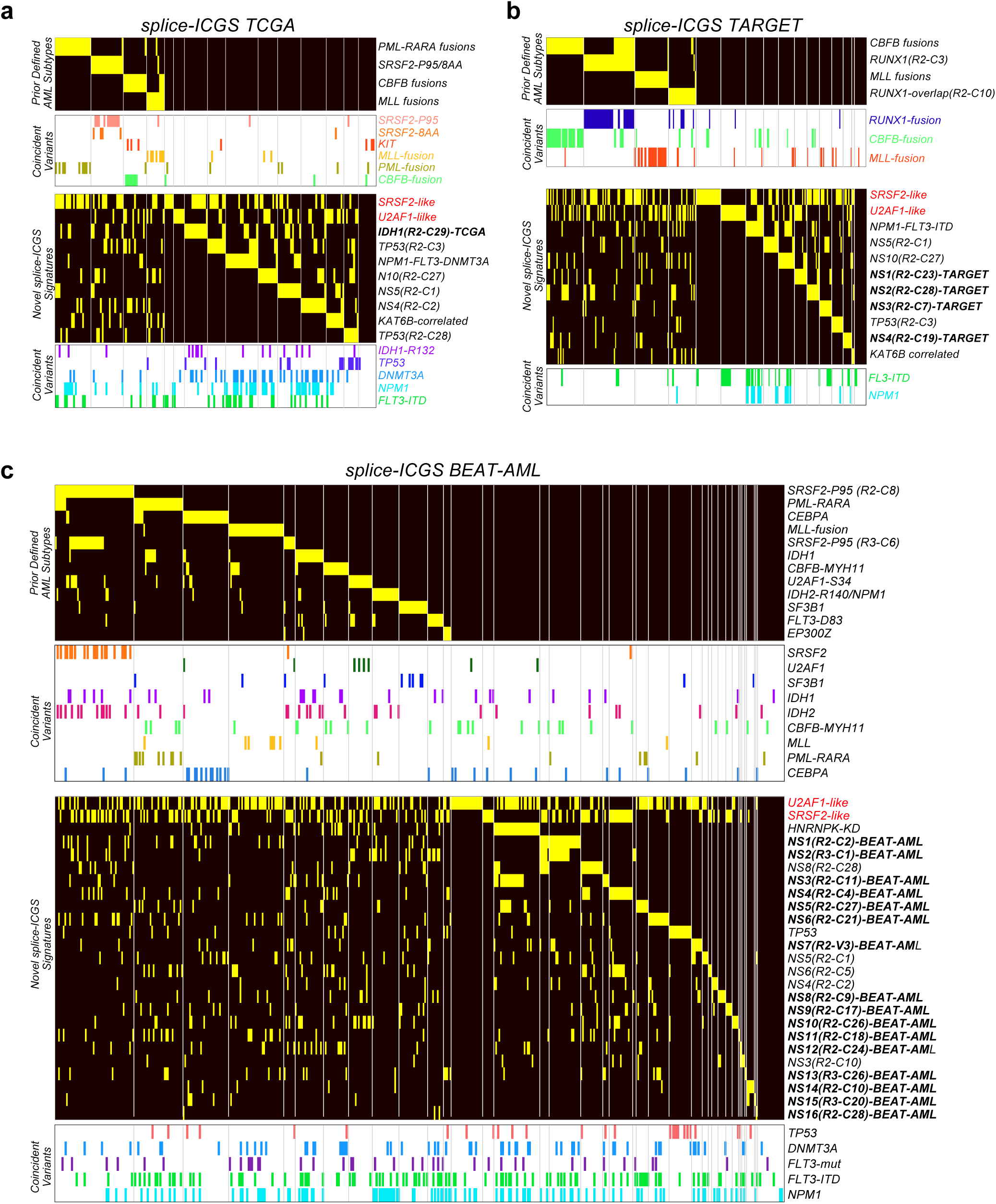
Reproducible AML subtypes identified in independent adult and pediatric AML cohorts. **a-c**) Splicing subtype predictions from splice-ICGS applied to TCGA (n=178),TARGET (n=257) and BEAT-AML (n=462). RNA-Seq samples. Subtype enriched mutations and labels are derived from OncoSplice (Supplemental Methods).

**ED Figure 3.**
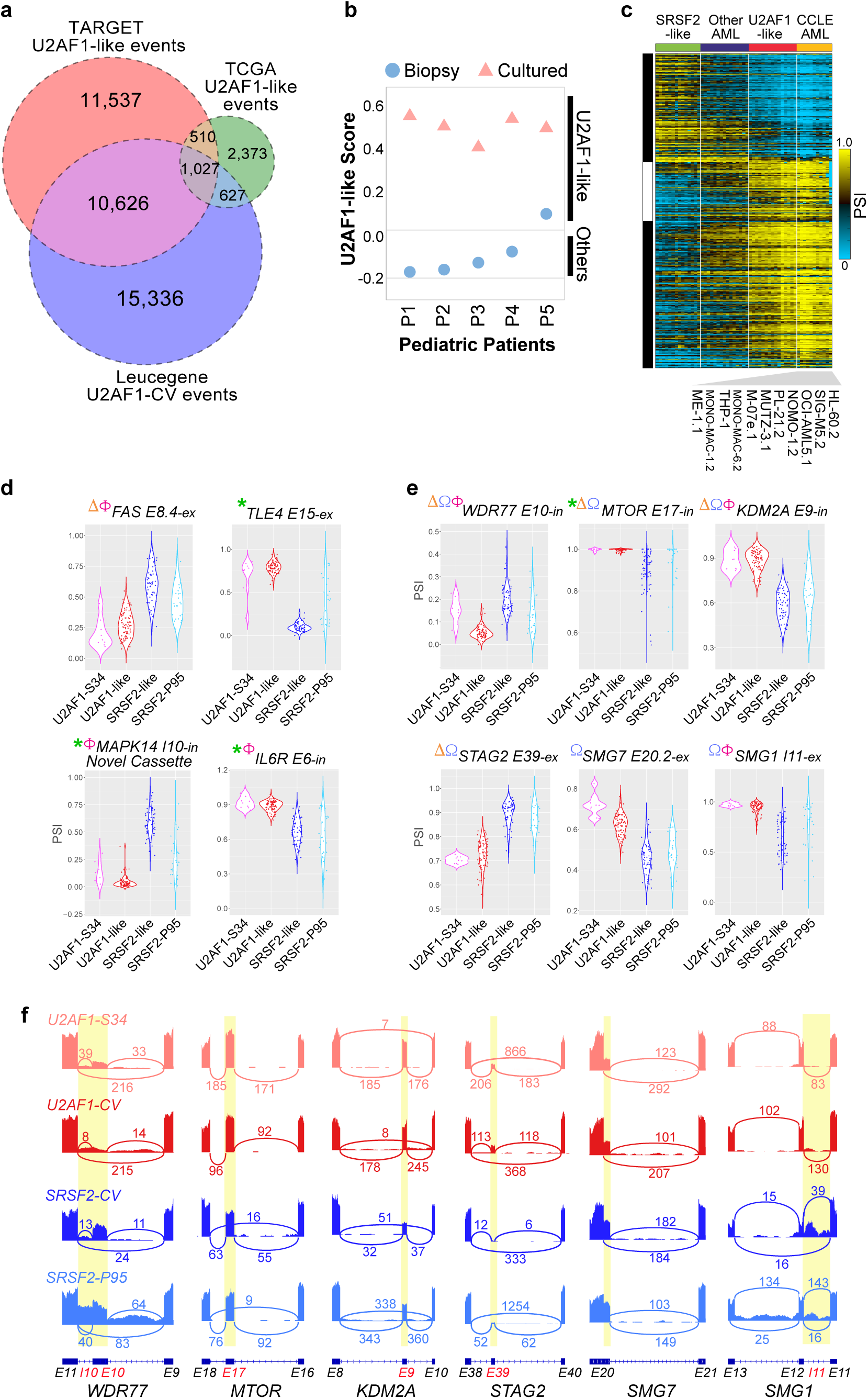
*U2AF1*-like splicing is associated with poor survival. **a**) Overlap (Venn diagram) of significant *U2AF1*-like alternative splicing events from distinct adult and pediatric AML datasets. **b**) *U2AF1*-like splicing status assigned to RNA-Seq data from primary pediatric AML patient bone marrow samples and matched cultured AML blasts. **c**) Heatmap of AML *U2AF1*-CV splice events in representative AML Leucegene samples compared to a panel of CCLE leukemia cell lines. **d,e**) Violin plots of selected *U2AF1*-CV splicing events in therapeutic cancer targets (d) and splicing event associated with poor overall survival in TCGA and TARGET (e). Splicing events predicted to result in loss of function (LOF) in *SRSF2*-CV (AltAnalyze and *in silico* translation) or genes that drop-out from the AML CRISPR screen (at least half of cell lines), are denoted with the indicated symbols. **f**) SashimiPlots for splicing evenets in panel e.

**ED Figure 4.**
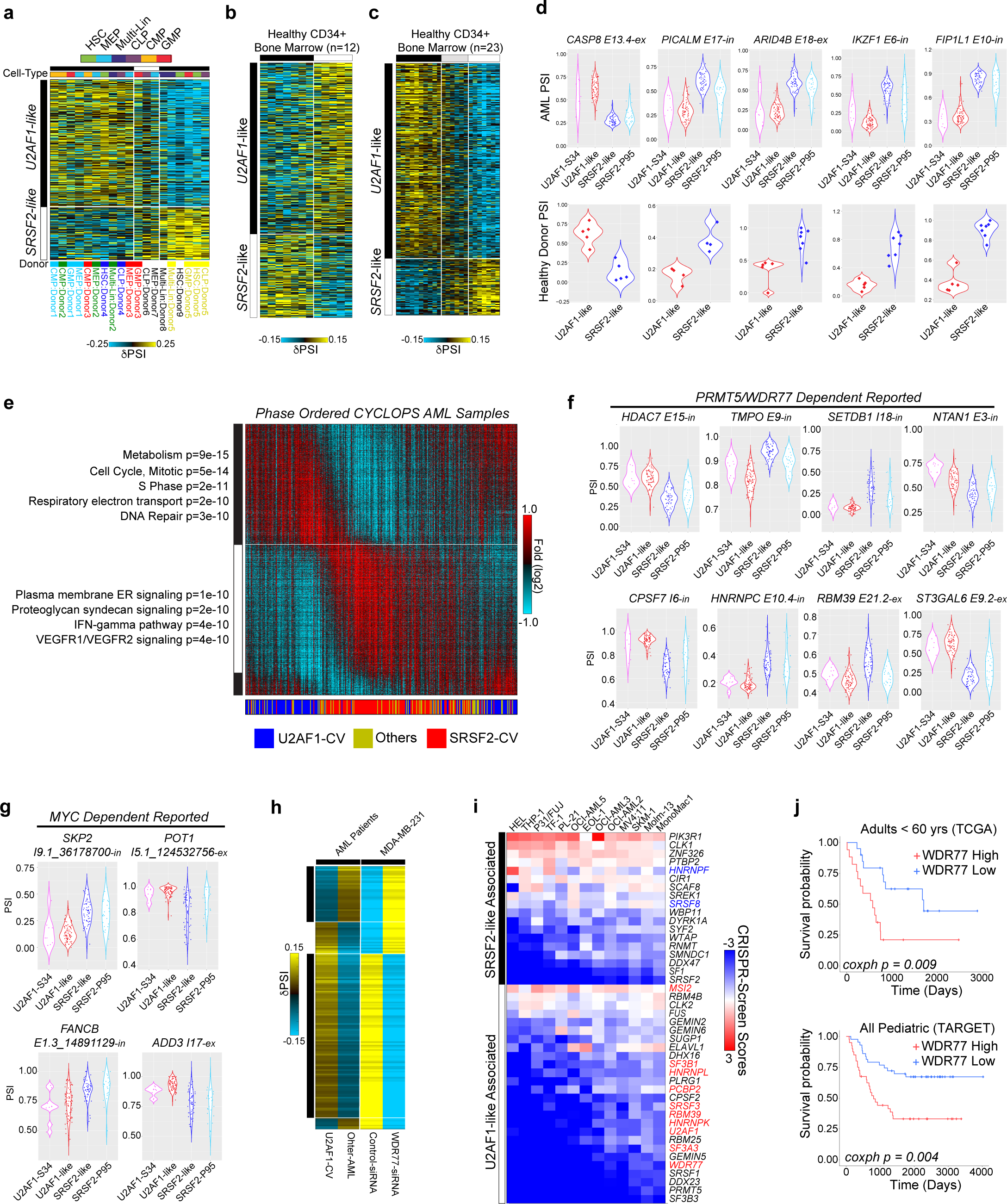
U2AF1-like predicts overall survival in adult and pediatric AML. **a**) Heatmap of the top 1,000 *U2AF1*-CV splicing events in sorted donor CD34+ progenitors (EGAD00001000745) indicates separation of samples (columns) by donor splicing pattern rather than capture gate (cell-types). **b**) Heatmap of 12 healthy-donor bone-marrow CD34+ cell RNA-Seq splicing profiles, HOPACH clustered based on the top differential 1,000 *U2AF1*-like AML splicing events. **c**) Heatmap of 23 healthy-donor bone-marrow CD34+ cell RNA-Seq splicing profiles (GSE111085). **d**) Violin plots displaying the PSI values for example *U2AF1*-like splicing events in AML (Leucegene) and healthy bone marrow progenitors of the 12 healthy donors. **e**) Heatmap of circadian ordered Leucegene AML samples according to estimated phase from the software CYCLOPS, filtered for genes with an FDR corrected p<0.5, relative amplitude >0.1 and R-squared >0.1 (ordered by acrophase). Top enriched Gene Ontology gene sets (GO-Elite) are shown on the left of the two circadian phase ordered gene clusters (GO-Elite Fisher Exact p-value shown). **f,g**) Violin plots displaying the PSI distribution for *U2AF1*-CV splicing events, previously reported to be regulated as a result of perturbation of PRMT5/WDR77 (f) or MYC (g). **h**) Heatmap of AML *U2AF1*-like significant splicing events overlapping with to shRNA of *WDR77* in MDA-MB-231 ^50^. **i**) Heatmap of individual AML cell-line CRISPR-screen scores (CSS) for RBPs associated with *U2AF1*-like or *SRSF2*-like splicing from gene expression or knockdown signature analyses. Red and blue labels indicate KD screen implicated RBPs whose expression is significantly correlated to either *U2AF1*-like or *SRSF2*-like splicing groups, respectively. **j**) Kaplan-Meier curves of WDR77 expression values 2±SD from mean in patients with AML from TCGA (top) or TARGET (bottom) with associated coxph p-values. Analysis of TCGA was restricted to cytogenetically normal AMLs with no RNA binding protein (RBP) mutations and under 60 years of age.

**ED Figure 5.**
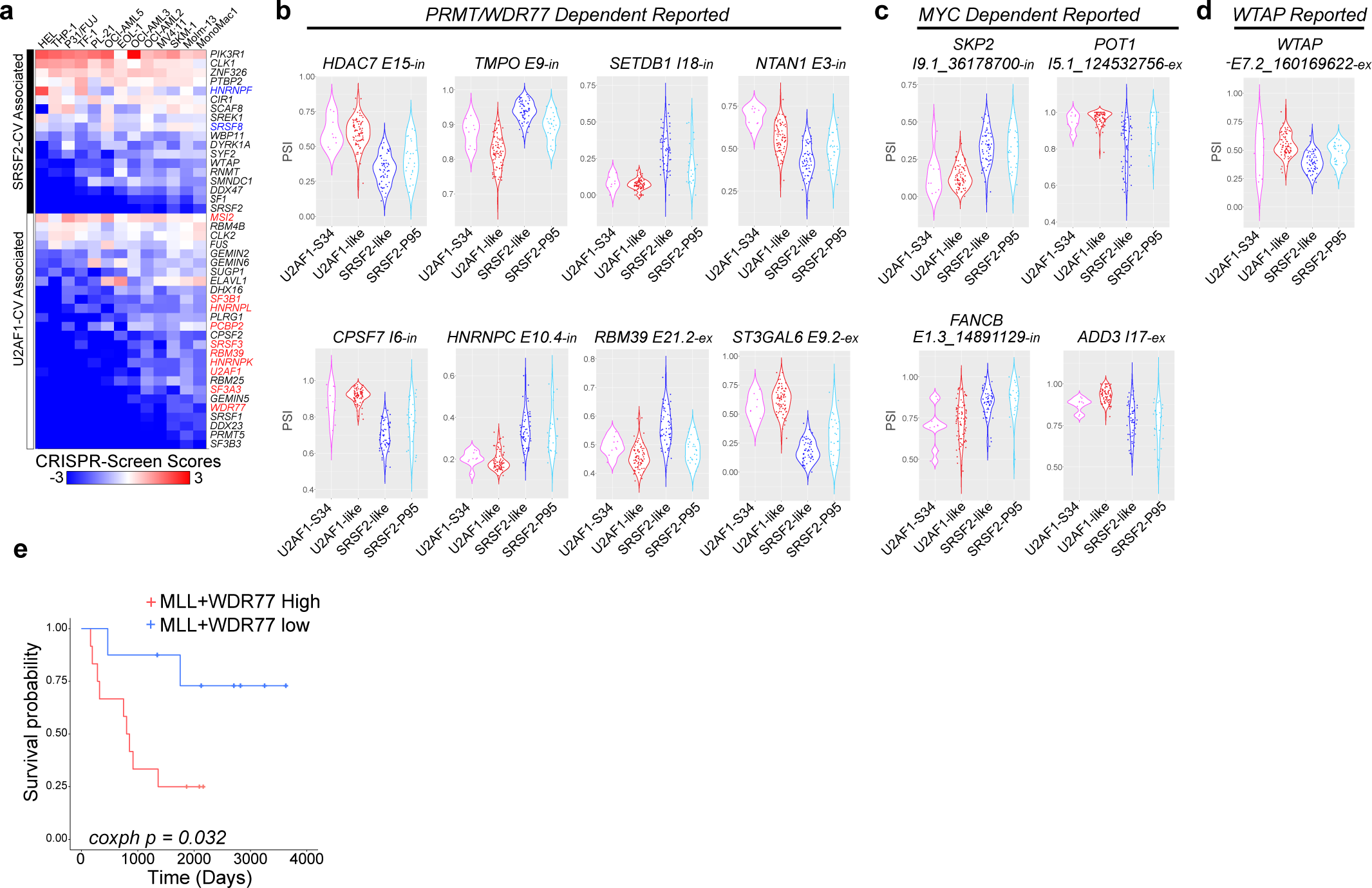

## Notes

https://www.synapse.org/#!Synapse:syn12103642

https://github.com/venkatmi/oncosplice

