## Supplementary Methods for "Broad de-regulated U2AF1 splicing is prognostic and augments leukemic transformation via protein arginine methyltransferase activation"

**SUPPLEMENTAL METHODS**

In the below sections, we detail the rationale and methods developed for the OncoSplice pipeline and associated cancer splicing analyses. Instructions for running the latest version of OncoSplice can be found at: <https://github.com/venkatmi/oncosplice>. A sample dataset is provided for analysis in the GitHub repository. Links to raw and processed data are provided in the online open-access Synapse database, where such data conforms to data sharing policy guidelines assigned to each respective dataset.

**Online datasets**: https://www.synapse.org/#!Synapse:syn12103642.

**Section 1: Unsupervised Identification of novel splicing-defined populations (splice-ICGS)**

1.1 – Technical Considerations for Splicing Subtype Detection

Although aberrant splicing is increasingly recognized as a driver in cancer, analyses of cancer associated splicing are almost exclusively limited to supervised analyses of known splicing factor mutations. Furthermore, while hundreds of cancer RNA-Seq datasets now exist, these datasets are rarely evaluated at the level of alternative splicing. Hence, to identify novel splicing subtypes in the absence of known causal mutations, we require accurate unsupervised splicing subtype discovery methods.

Various unsupervised gene expression subtype identification approaches have been described for cancer transcriptome profiling data. These include unsupervised clustering, PCA and bi-clustering approaches ^1-3^. However, such methods are not ideal for identifying novel alternative splicing patient subtypes from RNA-Seq, due to frequent missing values (data sparsity), variability in sequencing depth and redundant events detected within the same gene that will result in non-informative correlations. Furthermore, distinct cancer subtypes often overlap with each other, complicating precise detection. Short-read RNA-Seq (<150nt reads) is subject to uneven or partial transcript coverage resulting from inherent technical biases in the sample preparation, biological variation in gene expression and differences in sample sequencing depth. As a result, certain samples may not have reads mapping to the genomic positions where differential alternative splicing events were detected in other samples of the same dataset. These samples can thus be plagued by frequent missing values for specific splicing events, which can represent the absence of or non-measured expression. Depending on the sequencing depth of the experiment, we find that around 20-30% of the junction ratio counts are missing on average (data not shown). Furthermore, current clustering and bi-clustering techniques used for sample subtype identification in other biological measurement data such as gene expression are not designed to handle such a large fraction of missing values through imputation ^4^. Exon-exon junction based splicing detection methods such as the Percent Spliced In method (PSI) ^5^ produce splicing inclusion estimates (ranging from 0 – 1) that can have different distributions in datasets of different complexities and sequencing depths (unimodal, bimodal, multimodal or normal), making it difficult to apply models that only consider a Gaussian distribution. Another important aspect for unsupervised splicing detection is the dimensionality of the data. Typically, gene expression datasets are limited to 40,000 genes, but splicing datasets can contain tens to hundreds of thousands of splicing events, resulting in higher dimensionality with decreased sensitivity to identify low magnitude signal.

1.2 – Unsupervised Splicing Signature Detection with Splice-ICGS

To address these challenges, we designed a new computational workflow called OncoSplice that aims to identify and characterize complex disease molecular signatures at the level of alternative splicing (**Fig. 1a**). Recently, we described the development of a novel unsupervised gene expression population detection method for single-cell RNA-Seq (scRNA-Seq) called Iterative Clustering and Guide-gene Selection (ICGS) ^6^. ICGS identifies coherent gene signatures that are capable of distinguishing frequent, rare and transitional cell populations from scRNA-Seq gene expression datasets. Here, we extend the existing ICGS pipeline (splice-ICGS) to the analysis of bulk-RNA-Seq splicing profiles with variable missing data, patients with complex overlapping splicing signatures and biological/technological confounding effects. Within OncoSplice, splice-ICGS is used to identify the most variable splicing events present in a PSI-matrix file (MultiPath-PSI algorithm) and the most coherent splicing signatures through multiple rounds of signature identification and exclusion.

1.2.1 – Genetically-defined splicing archetypes in AML. Although AS has been previously implicated via splicing factor mutations in adult AML or translocation oncofusions in pediatric AML, wide-spread AS with genetically or non-genetically defined AML have not ^7^. Given that Leucegene has improved both the depth of sequencing and cohort size, our analyses focused on an assigned training (n=366) and test set (n=70, prior annotated set) from this dataset. To understand the mutational landscape of this incompletely characterized patient dataset, we determined commonly occurring and *de novo* point mutations, structural rearrangements, insertions and deletions in prior implicated cancer genomic loci (**ED Fig. 1a**). Compilation of predicted impactful variants (COSMIC database, literature) in RNA-binding proteins (RBPs) identified 79 patients (21%) in the training set with such variants (**ED Fig. 1b**). These included mutations in prior AML associated RBPs (*U2AF1, SRSF2, SF3B1, SF3A2, ZRSR2, PRPF40B, THRAP3*) in addition to rare co-occurring variants in *U2AF1*, *SRSF2* and *U2AF2*, not previously reported and accounting for 8% of patients with RBP mutations (ED **Table 1**). *SRSF2*-P95 mutations were found to be the most frequent RBP alteration (7%) in Leucegene as well as TCGA (8%) (which appear to have been missed in prior TCGA AML analyses ^8^).

1.2.2 – Splicing Event Detection with MultiPath-PSI. To consistently detect diverse alternative splicing and alternative promoter associated events, we adapted a recently developed splicing algorithm in the software AltAnalyze called MultiPath-PSI ^9-11^. This algorithm is introduced in AltAnalyze version 2.1.1 and requires aligned BAM files as input (inclusion of novel junctions and strand prediction are recommended). Similar to other recent reported local splicing variation (LSV)-based methods, such as LeafCutter ^13^, MultiPath-PSI considers the detected junctions within a restricted genomic interval for assessing local junction expression differences. Unlike these alternative approaches, MultiPath-PSI considers all known and novel exon-exon junctions in a sample or cell and computes its relative detection compared to the local background of all genomic overlapping junctions that can be directly associated with the given gene of interest (sharing at least one known gene-associated splice-site). This calculation provides a more inclusive and conservative estimate of LSV. These same junctions are used to identify high confidence intron retention splicing events evidenced by pairs of exon-intron and intron-only mapping paired-end reads, sufficiently detected at both ends of a given intron (5' and 3'). This more stringent algorithm requires that only counts for exon-intron spanning reads are reported, in which multiple exon-intron spanning reads are detected at both ends of the intron and a matching paired-end read contained entirely within the intron is present (BAMtoExonBed module of AltAnalyze). PSI values are calculated for any junction with a minimum PSI difference of 0.1 (10% PSI) between any samples or cells. PSI values are only reported for samples or cells for a given splicing event in which sufficient read-depth is present (minimum of 20 reads per examined junction interval). Junctions are clustered into groups of unique junction clusters when reporting the results to identify redundant splicing events. Unique junction cluster IDs are determined by examining the overlap in exon-exon and exon-intron junctions for a given gene to identify connected subgraphs in the exon-exon/exon-intron network. Differential splicing events can be calculated through this method using the associated metaDataAnalysis.py module in AltAnalyze for a set of supervised sample groups and comparisons of the input PSI values. This latter function further includes various splicing effect predictions including exon-inclusion or exclusion, redundant splicing-events (clusters), splicing-event type, predicted protein isoform differences and domain impacts (**Section 6**).

| 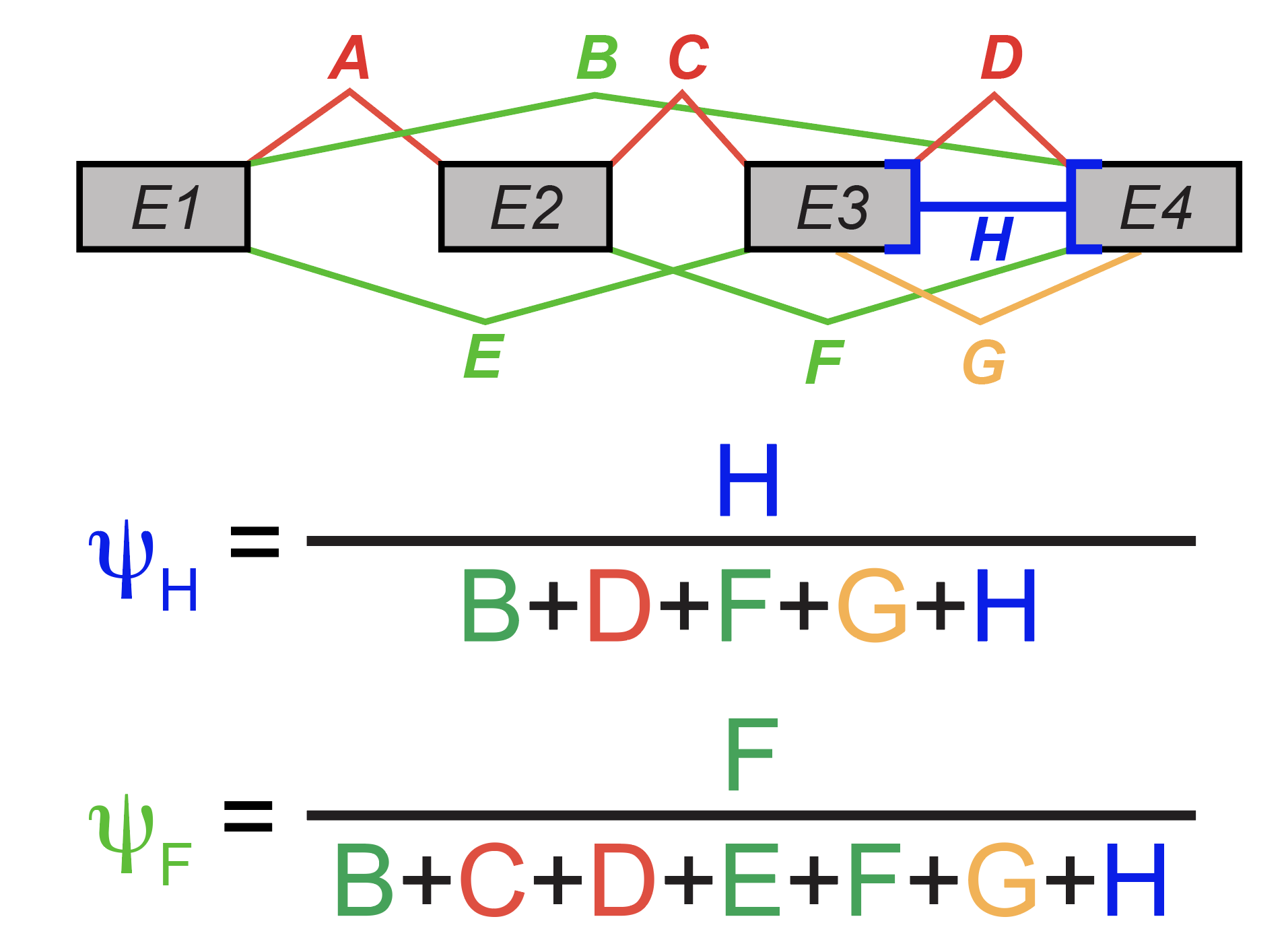 | MultiPath-PSI example splicing event calculation for intron retention (H) or more complex use cases (F), from exon-exon and exon-intron junction read count ratios. |
| --- | --- |

To identify an appropriate statistical method for comparison of splicing events between splice-ICGS identified cancer subtypes, we directly compared the results from multiple previously reported algorithms (Wilcoxon rank sum, Kolmogorov–Smirnov test (KS) and Empirical Bayes Moderated t-statistic [eBayes]) ^14-17^. We focused our comparison on well-studied cancer splicing subtypes (*SF3B1*, *U2AF1-*S34 and *SRSF2-*P95 mutations) compared to cytogenetically normal AMLs. Importantly, although PSI values were observed to be skewed towards the PSI extremes across the entire Leucegene dataset, splicing events themselves were normally distributed when comparing relative PSI values for individual splicing events (data not shown). For all the splicing-factor mutational comparisons, the Wilcoxon ranked sum and the eBayes tests produced consistent results with each, with the Wilcoxon overlapping almost entirely with eBayes predictions. The most consistent results (>92% overlap), were observed with *SRSF2-*P95, which has the largest number of splicing factor mutation patients. Importantly, several splicing events that were uniquely significant with eBayes were previously observed in an introduced cell-line *U2AF1-*S34 isogenic mutation, including events in the genes *MOK, ADARB1, MATR3, N4BP2, PDGFC, E2F6, TSEN2*, and *BRCA1* ^18^. As the eBayes method appeared to have excellent sensitivity, specificity and produce much faster runtimes, this method was selected as the default algorithm for all downstream splicing comparison analyses.

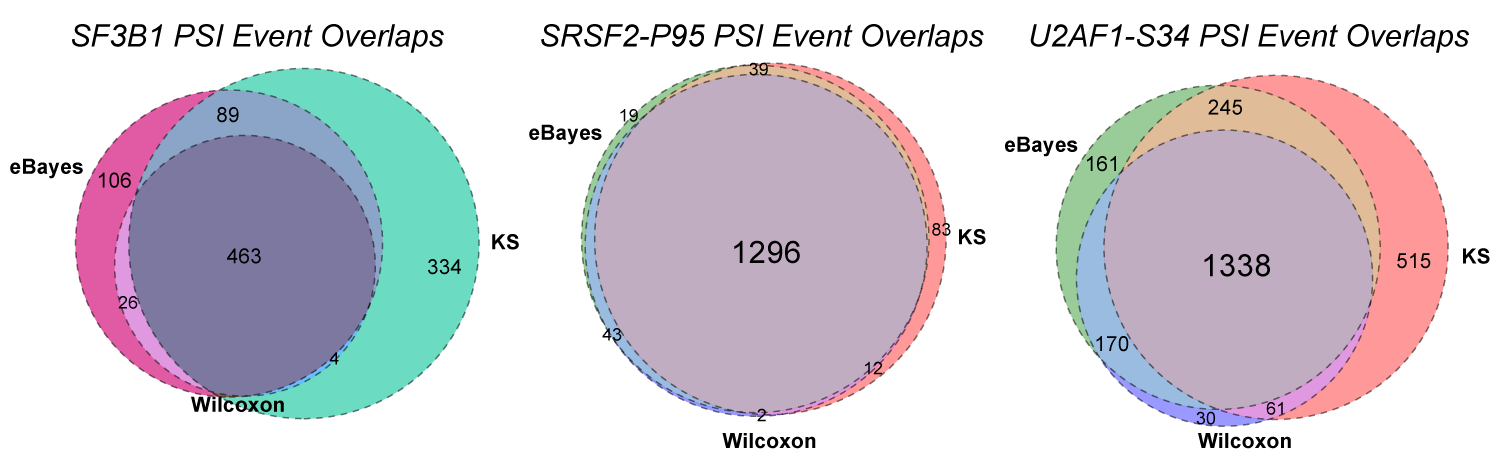

To assess the broad impact of AS in genetically defined subtypes of AML, we compared AS in AML with common structural rearrangements: *PML-RARA*, *CBFB-MYH11*, *RUNX1-RUNX1T1* and *MLL-*fusions. Using this algorithm on our training dataset, we find that oncofusion-specific splicing is a common feature of adult AML (**ED Fig. 1c**). Further, compared to gene expression, AS improves the overall detection of AML with oncofusions, based on a 3-fold cross validation analysis performed on the training dataset (patients with known fusions or mutations) (**ED Fig. 1d**,**e**). Such oncofusion-specific splicing was also verified in pediatric AML (**ED Fig. 1f**). For each of these oncofusions, 20-30% of these same splicing events were shared between adult and pediatric. Upon applying this supervised classification approach to the remaining samples in the dataset, using these unique splicing-event signatures we further discovered 6 novel patients with *SRSF2-8AA* deletions that were correctly classified as *SRSF2-P95* variants, but that were not detected in our initial variant analyses (**ED Table 1**). Although only previously confirmed in MDS and not in AML ^19^, this result is in agreement with prior studies showing that *SRSF2*-8AA and *SRSF2*-P95 mutations result in similar splicing impacts ^19^.

1.2.3 – Evaluation of MultiPath-PSI. To ensure that MultiPath-PSI is sufficiently robust to detect known and novel splicing events for downstream unsupervised analyses, we tested its ability to accurately identify experimentally validated alternative splicing events and benchmarked this method against alternative local splicing variation (LSV) tools (LeafCutter [version 0.2.7], splAdder [version 2.2.2] and rMATS [version 4.0.1]).

Experimental Validation: We compared MultiPath-PSI predictions to a previously reported RT-PCR dataset of 60 splicing events from mouse liver versus cerebellum ^20^. For this analysis, exon-exon junctions listed for each validated splicing event were matched to MultiPath-PSI reported junctions following alignment of the FASTQ files with STAR to mouse genome (mm10). All 60 splicing events were detected and considered differentially spliced (δPSI> 0.1, p<0.05, FDR adjusted) with MultiPath-PSI and had comparable changes in δPSI with the RT-PCR analysis.

| 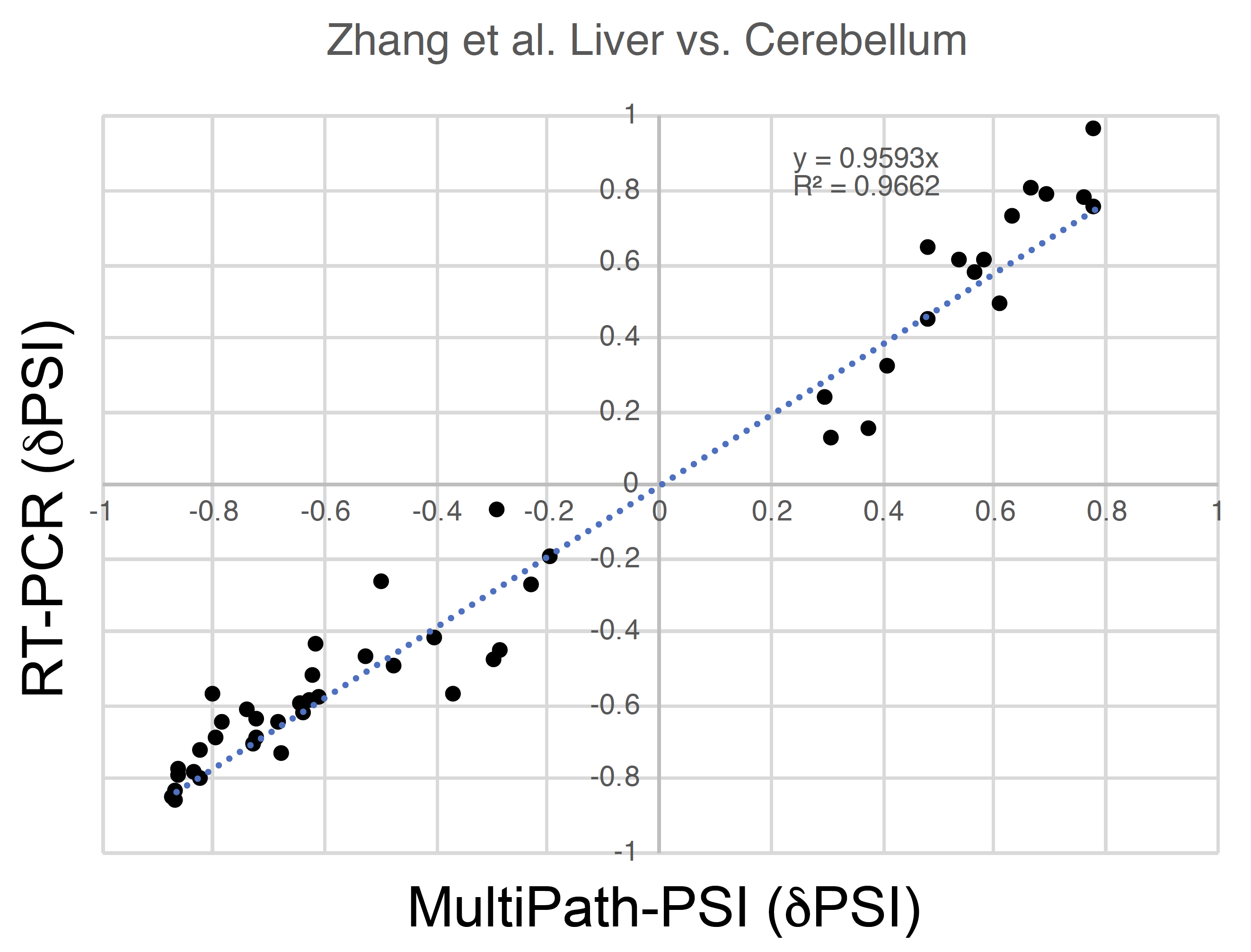 | **MultiPath-PSI accurately identifies experimentally validated tissue-specific splicing events**. Previously reported δPSI values for significantly differential RT-PCR splicing events were compared to MultiPath-PSI splicing events for 8 mouse liver and 8 mouse cerebellum samples, indicating high concordance. |
| --- | --- |

Synthetic Evaluation: As an unbiased evaluation of possible known and novel alternative splicing events, we developed a simulated RNA-Seq dataset derived from known and novel isoform predictions for pluripotent stem cells (PSC) and *in vitro* derived day 30 cardiomyocytes (CM) (https://www.synapse.org/#!Synapse:syn12104338). To create this simulation dataset, we first predicted known and novel isoforms and expression estimates using Cufflinks on biological triplicate samples (PSC and CM). To define benchmarking splicing events from this synthetic isoform data we created a python script to extract exon-exon and exon-intron junction pseudo-counts from all isoform files, computed PSI values for each junction relative to all overlapping junctions and finally obtained differential splicing results from those replicate PSI values. To accomplish this, we first determined the location of all introns from the Ensembl version 72 database and all annotated exon-exon junctions coordinates from any known isoform in Ensembl or UCSC genome database (AltAnalyze program: AltDatabase/ensembl/Hs/Hs_Ensembl-annotations.txt). Each isoform in the Cufflinks isoforms.fpkm_tracking file was then decomposed into its cognate exon-exon or exon-intron junction coordinates and assigned to a single Ensembl gene based on the genomic coordinates of each junction or evidence of retained intron/exon from the aforementioned exon/intron database using a custom python script (see below). Next, to obtain pseudo-counts for each junction, the FPKM values for all isoforms that contain the junction were summed and multiplied times ten and converted to integer values (see IsoformToJunctionCounts.py, https://www.synapse.org/#!Synapse:syn20486803). PSI splicing ratios were computed for each junction from the pseudo-junction counts, considering the counts for the evaluated junction (numerator) and the count sum of all genomic overlapping junctions including that junction itself, that correspond to the same gene (denominator). δPSI values were computed from the average PSI values in the PSC and CM groups along with p-values using limma (empirical Bayes moderated t-test p<0.05, Benjamini-Hocherg corrected). Overlapping splicing events were determined from a junction-graph of junction genomic coordinates for all significant benchmarking events (input and output files in the above Synapse link).

To compare the performance of each evaluated tool, we next produced simulated RNA-Seq database based on the original Cufflinks isoform abundance data. To do this, Cufflinks isoform GTF and expression values (isoforms.fpkm_tracking) were supplied as input for the software Polyester ^21^ to produce simulated RNA-Seq reads at a depth of 100 million paired-end reads using the software default parameters. These subsequent FASTA files were supplied as input for genome alignment (hg19) with the software STAR to produce BAM files with aligned, stranded, paired-end reads. The resulting aligned sequencing data were evaluated with each analysis method and benchmarked against isoform-derived alternative splicing events (exon-exon and exon-intron junctions) comparing CM to PSC using these input BAM files. For LeafCutter, splAdder and rMATS, the default recommended options were applied, including minimum junction count thresholds. Both MultiPath-PSI and LeafCutter report splicing-events clusters which assign overlapping splicing events to a common identifier. To identify matching splicing events in the three methods, exon-exon and exon-intron junctions were matched at the genomic coordinate level using a custom python script (comparisonAUPR.py) using one or multiple outputs from each of the evaluated tools. For rMATS and SplAdder, new unique splicing event clusters were obtained with this script, by creating a junction-graph of junction genomic coordinates from this method.

To evaluate the performance of the different algorithms in reporting the differential splicing events, precision-recall curves were generated using Matlab for the synthetic isoform splicing results (aupr_step.m script). For each method, the significant splicing events were ranked in descending order according to the empirical score or p-value reported by each algorithm. As rMATS does not detect novel splicing events and LeafCutter does not detect intron retention events, for each precision recall curve, only junctions that were detected in each program were considered in the ground state truth, which improved the overall results for these three methods (Filtered – see below table). The inputs, outputs and ReadMe file are provided in Synapse to clarify the processing and integration steps for each file. Please note, each algorithm reported only a portion of the benchmarking splicing events as significant. To generate precision-recall curves that extend to recall=1, the non-reported gold-standard splicing events were included additionally with a p-value=0 or lowest empirical score. This inclusion leads to the sudden peak in precision with recall identified in the tail of the precision-recall curves for each tool. Comparison of unique splice-junction clusters produced by each algorithm to these benchmarks indicated that MultiPath-PSI has the highest or comparable overall accuracy, based on precision and recall estimates **(Fig. 1c**). Hence, we expect MultiPath-PSI to be highly accurate for determining known and novel splicing events unsupervised splicing analysis (splice-ICGS).

| **Software** | **Sensitivity** | **All AUCPR** | **Filtered AUCPR** | **Synethic TP Events** | **Total Events** |
| --- | --- | --- | --- | --- | --- |
| *MultiPath-PSI* | 76.2% | 0.58 | 0.67 | 279 | 2257 |
| *Leafcutter* | 69.9% | 0.40 | 0.51 | 256 | 5138 |
| *SplAdder* | 70.5% | 0.16 | 0.31 | 258 | 6426 |
| *rMATS* | 39.3% | 0.10 | 0.21 | 144 | 4932 |

**Evaluation of MultiPath-PSI relative to other local splicing variation approaches using synthetic isoform splicing data**. AUCPR was calculated using junctions mutually detected by each specific application and the synthetic ground state truth (see Filtered AUCPR) or all synthetic junctions (All AUCPR). TP=true positive synthetic events. Note the term “events” in the table is used to represent unique splice-junction clusters instead of individual splicing events.

1.2.4 – Splice-ICGS. The core ICGS method performs expression filtering, pairwise correlation of dynamically expressed genes, iterative clustering and the selection of pattern-specific guide genes to identify highly coherent gene expression clusters. To handle the missing values from splicing PSI values, pairwise correlations are calculated using only the indices with values present in both of the splicing events, through a remote call to the R built-in correlation function. Only correlations with a Pearson coefficient greater than the user supplied threshold (by default rho>0.4) and a correlation p-value <0.05, occurring between distinct genes (non-redundant splicing events) are retained for further analysis. Similar to the conventional ICGS method, PSI and sample clustering is performed using the method hierarchically ordered partitioning and collapsing hybrid (HOPACH) through a remote call from Python, accounting for missing values (hopach package in R). As in ICGS, one splicing-event is selected from each identified splicing-signature (splicing-event cluster) and used for supervised selection of additional splicing events from the initially filtered PSI value file. This analysis is designed to identify the most coherent and correlated splicing patterns. Following the first round of splice-ICGS, the results from this analysis (Guide-3 selected splicing signatures) are subjected to two separate analyses: 1) the predominant splicing signatures are selected as potential biological and technological confounding effects and eliminated from the dataset for subsequent rounds of splice-ICGS (Signature Depletion - **Section 1.4**) and 2) a more rigorous NMF-based method for subtype identification analysis is applied to the samples from the current round of results (Subtype Identification - **Section 1.3**).

1.3 – Splice-ICGS Sample Subtype Classification.

Typical unsupervised population discovery methods identify discrete groups of non-overlapping samples. While sufficient for non-complex disease subtypes or single-cell populations, such methods are unable to distinguish molecular signatures arising from samples with complex disease genetics and clonal or cellular admixtures. For example, splicing factor mutations often co-occur with gene-fusions (e.g., MLL-fusion and *U2AF1-*S34 mutations) ^22^. Several techniques exist that perform feature extraction, such as PCA, ICA, non-negative matrix factorization (NMF) and SVD. Following the core-ICGS analysis, the workflow applies sparse NMF to non-negative PSI values to delineate distinct splicing signatures followed by preliminary subtype splicing event detection and subsequent supervised patient sample classification (**Fig. 1d**).

1.3.1 – Sparse Non Negative Matrix Factorization (Sparse-NMF)

Following the ICGS step, splice-ICGS applies sparse NMF to non-negative PSI values to delineate distinct splicing signatures followed by preliminary subtype splicing event detection and subsequent supervised patient sample classification. HOPACH clustering of samples is challenged by the identification of subtypes given overlapping splicing event signature. Hence, we turned to sparse NMF as a possible downstream method to refine subtype detection. Given a non-negative matrix *A* of *mxn* size, where each column of *A* corresponds to a data point in the *m*-dimensional space, and a positive integer, NMF finds two non-negative matrices and so that *A~ SG*, where S~R^mxk^ is a basis matrix, G~R^kxn^ is a coefficient matrix. Due to k<m, dimension reduction is achieved and a lower dimensional representation of *A* in a *k*-dimensional space is given by *H*. For improved cluster identification, Sparse-NMF (SNMF) was introduced. SNMF uses a L1-norm minimization and is solved using a fast nonnegativity constrained least squares algorithm (FCNNLS) ^23^.

For PSI splicing analyses, A is an *mxn* matrix, where m is samples and n is the splicing event (transposed ICGS Guide3 matrix). Optimization solution for S, G is given by:

$$\min_{W,H} f\left( S,G \right)=\frac{1}{2}{|\left| A-SG \right||}_{F}^{2},s.t. S,G\geq0,$$

Additionally, sparsity constraints are imposed on the S and G matrices using L2 and L1 norm respectively,

$$\min_{W,H} \frac{1}{2}{|\left| A-SG \right||}_{F}^{2},+ \eta{|\left| S \right||}_{F}^{2}+\beta\sum_{j=1}^{n} {|\left| G\left( ;,j \right) \right||}_{1}^{2}s.t. S,G\geq0,$$

Where 𝛈>0 is a parameter that is used to suppress ||S||_F_ and 𝛃 >0 that is trade-off between accuracy and sparseness.

To identify sample clusters, this workflow uses a sparse NMF algorithm that controls the degree of sparseness in the coefficient matrix *H (*splicing events) via alternating non-negativity-constrained least squares. To perform NMF analyses, the software must select the number of ranks (patient subtypes) to be used. The results from splice-ICGS are used to calculate this rank. Splice-ICGS uses HOPACH clustering to perform clustering on the splicing events, however, clusters that are highly correlated to each can be subdivided sometimes into multiple clusters, thus to improve the cluster detection, intracorrelation is performed among all the splicing events to aggregate highly similar HOPACH subclusters. The resultant clusters are used by NMF to estimate the ranks. If the number of clusters=2, then NMF rank=2 (implies a confounding broad splicing signature) and if number of clusters >2, rank=30 (NMF default). The goal of setting rank=30 is to overestimate the rank and consider only subtypes that are validated (have significant number of differential splicing events). Clusters without significant differential splicing events are discarded from the analysis, reducing the final rank to a much smaller, but optimized value. We refer to this step as “cluster fitness” (only the fittest clusters survive). Upon running NMF, a reduced basis matrix is returned whose dimensions are n x k, where n is the number of samples and k is the number of ranks provided. To identify sample clusters, the resultant basis matrix is binarized. We use snmf function provided in the nimfa package.

1.3.2 – Assigning NMF Subtype Labels to Individual Samples. For each splice-ICGS subtype, a value = 1 is assigned to samples with an NMF score > mean+2std of all NMF scores for that subtype. The samples with a value = 1 are considered representatives of each subtype of k. Thus, a single sample can be assigned to multiple clusters. For each of the clusters, splicing events that were differential spliced (adjusted p-value<0.05 and dPSI >|0.1|) were identified using the metaDataAnalysis.py module in AltAnalyze. To find valid and cohesive sample clusters two criterions were introduced:

1) Clusters with <100 differential splicing events are removed, since such a cluster can be considered an artifact of NMF.

2) Given that the same samples can be assigned to multiple clusters in each round, cluster redundancy is likely to occur. Such clusters may represent finer granularity within a larger cluster or may be largely redundant. For smaller cluster(s) embedded within a larger sparse-NMF cluster, if over 50% of the samples overlap with >70% overlap in the specific splicing events associated with those clusters, such sub-clusters are considered to be redundant with the larger parent cluster and hence are not reported as distinct subtypes.

For each of the remaining subtypes, a centroid is computed by taking the average of the specific representatives of the subtype across all the differentially spliced events. These centroids and alternative splicing events are used as training data for a One vs Rest Linear SVM classifier which then is applied to the entire dataset to re-classify samples for each subtype. The one vs rest svm classifier fits one classifier for every subtype vs remaining subtypes ^24^. Given a set of training data (*x_1_,y_1_*),(*x_2_,y_2_*)…(*x_n_,y_n_*) where *x_i_*~R^n^, i=1,…*.n* and y~{1,….*k*} where *k* is the number of classes. The *l^th^* SVM classifier solves for:

$$\min_{w^{l},b^{l},\xi^{l}} \frac{1}{2}({w^{l})}^{T}w^{l}+C\sum_{i=1}^{l} \xi^{l}$$

$({w^{l})}^{T}\varphi(x_{i})+b^{l} \geq1-\xi^{l} if y_{i}=l$

$({w^{l})}^{T}\varphi(x_{i})+b^{l} \leq-1+\xi^{l} if y_{i}\neq l$

$\xi^{l}\geq0, i=1,\ldots..n$

where φ is a function on the training data and $C\sum_{i=1}^{l} \xi^{l}$ is a penalty parameter.

Solving this *gives* *k* decision functions:

*d_1_*=(${w^{1})}^{T}\varphi(x)+b^{1}$,

*d_k_*=(${w^{k})}^{T}\varphi(x)+b^{k}$

Instead of using the largest value across the *k* decision function to decide the class, we consider

`	x_i,l_ =1 if $d_{i,l}$> 0

x_i,l_= 0 if	$d_{i,l}$≤ 0 where i=1,…*.n* and l = 1…*k*

Thus, while a typical SVM will assign a sample to a single subtype with the highest decision function score, our approach assigns a sample to any subtype with a decision function score greater than zero.

This schema allows for samples to be classified to multiple subtypes or no subtype. When NMF is run with rank=2, rather than automatically assigning all patient samples in the dataset to one of the two groups, the SVM will output a stringent (decision function score > 0.5 or <-0.5) and an alternative conservative (aka “expanded”) (decision function score > 0.25 or <-0.25) estimate of which samples are assigned to which group based on the decision function score. For this, we call the linearSVC function provided in scikit-learn.

1.3.3 Evaluation of splice-ICGS compared to existing clustering and Biclustering approaches. To assess the relative performance of splice-ICGS in detecting informative splicing-defined subtypes, we analyzed known AML splicing-defined subtypes using to a broad range of established unsupervised clustering methods. While no dedicated unsupervised splicing approaches exist to our knowledge, we selected algorithms for comparison that have been deemed potential gold standard unsupervised subtype detection approaches for high-throughput genomics datasets: PLAID ^25^ (iteratively identifies overlapping confounding signatures), EBIC ^26^ (identifies trend-preserving, overlapping subtypes for both narrow and broad biclusters), QUBIC2 ^27^ (biclustering approach recently developed for RNA-Seq data), RuniBIC ^28^ (identifies overlapping biclusters) and FABIA ^29^ (identifies overlapping biclusters in RNA-Seq data). We additionally tested two reported state-of-the art methods for single-cell RNA-Seq (scRNA-Seq) cluster determination: Seurat 2.3.4 ^30^ and SC3 ^31^. Although, such scRNA-Seq algorithms have not been previously evaluated for the evaluation of bulk RNA-Seq or alternative splicing, these are highly refined, multi-step dimensionality reduction approaches designed to find rare and common (but not overlapping) clusters in highly skewed high-dimensional datasets. The relative accuracy of each method to identify pre-defined AML splicing subtypes was evaluated using an F1-score (harmonic mean of precision and recall). Since, single cell unsupervised techniques are applied to counts data and cannot handle missing values, we initially filtered the dataset for splicing events with >75% values (~88,144 splicing events) and then imputed the remaining missing values with the median splicing event value calculated for the specific splicing event across all samples for all methods except splice-ICGS. The resultant PSI values were multiplied by 1,000 to resemble counts dataset. All the biclustering (EBIC, QUBIC2, RuniBIC, FABIA, PLAID) and clustering (Fuzzy-C, GMM) approaches were run with default parameters with k set to 9 (true positive clusters=9). Of these approaches, only PLAID and EBIC detected at least two of the 9 subtypes with high sensitivity and specificity. Both these approaches detected the *SRSF2*-P95 subtype with high sensitivity. However, these approaches failed to detect the oncofusion subtypes and the smaller splicing factor mutation subtypes. To run Seurat, we adjusted the filtercells function parameter based on the QC plots. The number of PCs considered for the FindClusters function was based on elbow plot (PCs=20). At a resolution of 3.0, Seurat v2.3.4 identified 23 clusters and we evaluated the performance of the tool in detecting the major mutation defined subtypes of AML and oncofusions. Seurat gave two clusters when run with these parameters. We varied the resolution parameter between 0.5-4 to further increase the number of clusters identified by the tool. The clustering and biclustering approaches where run with k=50. As shown in ED Figure 1i, alternative unsupervised methods fail to identify subtypes with few samples even though they are associated with a strong splicing signature (narrow biclusters). Of the 9 subtypes examined, Seurat only identified the *SRSF2*-P95 mutation with high specificity and *RUNX1* and *CBFB* fusions were identified as a single cluster. For SC3, we used the default values for the filtering step but varied the range of k’s (number of clusters) between 2-30. For each cluster size, SC3 provides the average silhouette width, stability and the number of differential events. We considered the cluster size (k) with the highest average silhouette which was 17 for the datasets. SC3 performed well in identifying 4 of the 5 onco-fusion subtypes (except MLL) and the *SRSF2*-P95 mutation cluster. However, it could not detect any of the 4 smaller splicing factor mutation subtypes with high precision and recall. Hence, splice-ICGS substantially improved the detection genetically defined AML splicing subtypes.

1.4 – Splicing Signature Depletion and Subsequent splice-ICGS Rounds.

Biologically confounding splicing signatures that represent a predominant signal in the data are excluded by splice-ICGS to improve the sensitivity to detect minor subtypes (e.g., splicing-factor mutation associated). This step of splice-ICGS is used to exclude (deplete) the identified subtype associated splicing events from the dataset and generate the input for the next round of the splice-ICGS, until the number of clusters for the rank = 1. To remove these signatures from the data, we use the actual values from the NMF w matrix for the selected subtypes with valid sample clusters and remove the splicing events correlated (rho <0.3) with the subtypes. This step improves the detection of minor subtypes that are obscured by major subtypes identified in the prior ICGS rounds. This method is implemented in the OncoSplice python module Correlationdepletion.py.

1.5 – k-means Clustering.

In the final step of the analysis, splice-ICGS uses k-means rather than NMF/SVM to identify the final remaining subtype. K-means clustering is performed instead of sparse NMF on two conditions: 1) when rank =1 and 2) when the number of subtypes identified with at least 100 alternative splicing events =1. This clustering considers k=2 and defines a new subtype based on the number of samples (class with the fewest samples) and whether it has a significant number of differential splicing events.

1.6 – Variant and External Subtype Enrichment Analysis of Splice-ICGS Clusters.

Once splice-ICGS has completed its assignment of patients to novel subtypes, the program must associate these patient subtypes with known metadata. Typically, variants include prior associated cancer mutations, insertions or structural/gene rearrangements but they can also include SNPs or other subtypes (e.g., FAB classifications), as we have done here. This metadata is supplied to the MutationEnrichment.py OncoSplice module in the form of a tab-delimited text file with patient sample ID and variant/subtype notation information (example inputs can be found at: <https://www.synapse.org/#!Synapse:syn12104087>). The input for subtype enrichment analysis is the final splice-ICGS NMF binary sample output file, as shown in the above link. This module performs a z-score and Fishers Exact Test enrichment analysis to provide p-values for the associations of this metadata with patient subtypes.

1.7 – Supervised Classification of Patient Samples from Other Cancer Datasets from Existing Splice-ICGS Subtypes.

In addition to the binary classification of patients from the primary analyzed RNA-Seq dataset into splice-ICGS subtypes (**Section 1.3**), we devised a method to identify the same likely subtypes in other external cancer RNA-Seq datasets or test samples from the same dataset. While test samples can be directly matched to the splicing PSI profiles from the same dataset, the same analysis becomes challenging when classifying subtypes from independent datasets processed in different laboratories, with different sequencing depth, library preparation methods and with different technologies (e.g., Illumina GAII for TCGA). To overcome this challenge, we first identified previously described patients with RBP mutations in the TCGA AML RNA-Seq dataset (e.g., *SF3B1, U2AF1-S34F, U2AF1-Q157*) or from splice-ICGS (*SRSF2*-P95) and tested prediction accuracy using various machine learning classification algorithms with Leucegene splicing profiles as references. Ultimately, we selected the One vs. One multi-class SVM classifier. Only unique splicing events are used for this analysis (unique event cluster ID), comparing the δPSI values (group PSI – average PSI of the two groups) between the splicing-subtype of interest versus all compared samples. The use of δSPI values is likely to be dependent on the exact PSI estimates from an individual experiment, which may vary for different library preparation and sequencing methods. This algorithm accurately classified all of the prior annotated splicing factor mutant patients in the TCGA AML dataset (n=6), including *SRSF2*-P95, *SRSF2*-8AA deletion and *HNRNPK* miscellaneous variants, not previously annotated in TCGA (**ED Table 1**).

**Section 2: RBP Signature Correlation Analysis (Bridger)**

To augment the set of unsupervised splicing-defined subtypes from Splice-ICGS, we developed a series of supervised methods: 1) to identify splice-ICGS or other putative novel subtypes associated with the gene expression levels of individual RBPs (Section 2.1), and 2) to identify patients whose splicing profile matches that of an RBP KD signature (Section 2.2).

2.1 – RBP-PSI Correlation Analysis.

To further identify novel splicing-defined subtypes from cancer RNA-Seq data or identify novel RBPs associated with splice-ICGS subtypes, we integrated an RBP-splicing event correlation method into the OncoSplice workflow. A large dataset of putative RBPs was collected using the Gene ontology and augmented based on the literature. The SpliceEnricher.py module of this workflow then calculates the correlation (Pearson) of the log2 expression of gene TPM or RPKM values relative to all splicing PSI values in the dataset using missing value correlations and a minimum of 20 non-missing PSI values (20 patient samples) for a give splicing event. The algorithm then reports RBP/splicing-events that are associated using a user supplied correlation threshold (rho=0.4 by default). Finally, the algorithm compares a directory of MultiPath-PSI splicing results (splice-ICGS subtypes) to identify RBP associated splicing signatures enriched in a particular subtype (Fishers Exact Test p-value). The result is a tab-delimited text file with RBPs and associated enrichments statistics. For the Leucegene dataset, one novel splice-ICGS signature was associated with *KAT6B* gene expression (*KAT6B* annotated subtype). Although *KAT6B* (aka *MORF*) oncofusions have been described in pediatric AML and other cancers ^32,33^, we were not able to identify any such fusions in our *KAT6B* high splicing-subtype.

2.2 – RBP KD Signature Classification Analysis.

Various genetic alterations in cancer, such as gene rearrangements, segmental duplications, amplifications, gene deletions and other cytogenetic abnormalities, result in the gain or loss of oncogenes that can impact downstream pathways necessary for oncogenic transformation or therapy resistance. We surmised that disruption or duplication of RBPs associated with distinct splicing subtypes, independent of the number of patients, may be better detected using existing KD RNA-Seq data. Hence, we integrated an additional component into OncoSplice to supervise the classification of patient subtypes from KD PSI signatures. To obtain the signatures, we analyzed ENCODE RBP and Lymphoblast KD (GSE52834) profiles following MultiPath-PSI analysis using the same supervised sample classification framework applied in Section 1.6 to all PSI datasets (non-adjusted eBayes p<0.05, δPSI>0.1). Again, we excluded the *U2AF1-*LIKE splicing signature as a potentially confounding variable for these analyses. For our analyses, this workflow identified a single subset of patients as *HNRNPK* KD associated in Leucegene, TCGA and TARGET. These predictions were confirmed between the Lymphoblast KD and ENCODE *HNRNPK* KDs. Downstream analysis of *HNRNPK* KD signature samples revealed that patients could be associated with a single nucleotide *HNRNPK* insertion, deletion, partial chromosome loss or *HNRNPK* novel intron retention using only this splicing signature. In addition to these, both *KAT6B* and *MLL* subtypes were frequently classified as *ACIN1* KD, while *SRSF2*-P95 patients were frequently classified as *PTBP1* KD.

**Section 3: RBP Prediction Analysis (RBP-Finder)**

To evaluate the potential contribution of RNA-binding proteins (RBPs) to the direct or indirect regulation of splicing for observed cancer splicing-defined subtypes, we developed a multi-parameter RBP enrichment prediction workflow called RBP-Finder. This workflow combines three distinct analyses: 1) CLIP-Seq enrichment of differentially spliced exons, 2) RNA-recognition element (RRE) enrichment and 3) RBP gene expression. Independent statistical enrichment analyses are performed within each step in this workflow and then relative scores are combined to arrive at a final weighted score representing the likelihood that a given RBP is involved in a particular splicing event.

3.1 CLIP-Seq Subtype Enrichment Analysis.

CLIP-Seq data provide experimental evidence of RBP RNA binding locations in the transcriptome in a specific cell type and can hence indicate regulation of splicing as well as other possible mechanistic impacts. We developed a method to compare spliced exonic/intronic regions to experimentally observed RBP binding locations that is based on a modification of the RELI method for intersecting genetic variants with ChIP-seq datasets ^34^. For these analyses, we assembled 180 CLIP-Seq datasets with peaks stored in the bed file format deposited in GEO or available from recent RNA ENCODE consortium efforts (Online Methods). To determine likely RBP regulators for specific identified cancer subtypes, we queried splice-ICGS differentially spliced exonic or intronic regions for patients versus all other AML cohort samples, following *U2AF1-*LIKE splicing signature depletion (Section 1.4). For each splicing event, MultiPath-PSI predicted regulated exonic or intronic regions (including +500nt and -500nt flanking sequence) were stored to a genome coordinate bed file. Each of these input bed files was then intersected with the library of CLIP-Seq peaks in order to identify splicing events that may be controlled by the binding of the corresponding RBP. An enrichment analysis was performed by permuting the location of CLIP-Seq peaks throughout the transcriptome to determine the likelihood that the number of RBP peaks associated with the splicing signature is likely to occur by chance alone. A total of 2,000 permutations were performed for each CLIP-Seq dataset, and the observed intersection counts were compared to the resulting distribution using a standard Z-score transformation. The Z-score was converted to a p-value and Bonferroni-corrected, producing a tab-delimited text file for each splicing signature/CLIP-seq dataset pair, which was used for integrative RBP scoring (Section 2.3).

3.2 RRE Subtype Enrichment Analysis.

In addition to the CLIP-Seq enrichment analysis, an RRE enrichment analysis was performed using the same exonic/intronic queried regions as input. These sequences were scanned using a large database of RRE motif models in the form of position frequency matrices (PFMs) taken from the CisBP-RNA database ^35^. These models were used for motif enrichment analysis using the HOMER software package ^36^. HOMER produces a series of HTML files, which were parsed to extract p-values for integrative RBP scoring (Section 2.3).

3.3 RBP Subtype Prediction Analysis.

Different RBP analyses will often implicate distinct possible splicing regulators; however, in many cases we have prior knowledge indicating which particular RBPs are responsible for a splicing subtype (e.g., RBP mutation, deletion). To optimize the identification of valid RBPs for each subtype, we designed an integrative model that considers multiple pieces of regulatory evidence. Estimation of the weight for each of these analyses in the overall cumulative score was performed using a logistic regression model. Logistic regression models are generally used to find the association of a set of independent variables with a binary outcome. In our case, the independent variables are the resultant p-values obtained from 1) RELI CLIP-Seq enrichment, 2) HOMER RNA recognition element binding site enrichment and 3) RBP up or down-regulation differential expression (subtype-included samples versus all other samples, eBayes t-test p-value, FDR corrected). To train the logistic model, two signatures with known RBP regulators were considered (*HNRNPK* loss of function and *SRSF2*-P95 mutation). These two factors were selected for training, as they represent diverse modes of RBP regulation (loss of function versus altered binding site preference, respectively). For these analyses, the p-values were converted to z-scores and for each signature, the binary outcome was assigned 1 for the expected RBP and 0 for the remaining RBPs. The model was trained and the coefficients of each of the analyses were obtained and applied to the remaining data. For each of the RBPs in the remaining signatures, a score was assigned by aggregating the product of the coefficients of the analyses to produce the final z-scores for the RBP. The top 5 RBPs with the highest score (>0) in each signature were considered as the top candidate regulators for that signature.

OncoSplice identified 11 splicing factor mutation specific signatures and 14 signatures with unknown splicing factor regulators. Of the 11 signatures with known splicing regulators, six had motif and CLIP-Seq information. RBP-Finder identified the correct splicing factor for five (*U2AF1*-S34, *SRSF2*-P95, *SRSF2*-8AA, *SF3B1* and *HNRNPK*) of these six signatures among its top 5 predictions (failing for *U2AF1*-Q157) . Note that *HNRNPK* and *SRSF2*-P95 were used for training the model and hence are inherently biased towards these predictions. As an independent evaluation of the RBP-Finder predictions for the other 14 splice-ICGS signatures (oncofusions and novel), we evaluated 232 RBP-KD PSI profiles using RNA-Seq data from ENCODE and GEO (see Online Methods). All 14 signatures had at least one RBP-Finder predicted regulatory RBP with an associated KD splicing profile among the top 5 RBP-Finder hits. In 8 out of the 14 splice-ICGS signatures (57%), we observed independent evidence for at least one RBP with concordant or discordant (>70% overlapping events changed in the same direction or opposite) among the top five hits, providing strong support for these RBP-Finder predictions (**ED Table 3**). Predictions from this logistic regression model were compared to those from CLIP-Seq, RRE (HOMER) and gene expression alone. Based on the p-values obtained from these approaches, the top five enriched RBP were selected and evaluated for at least one RBP with concordant or discordant (>70% overlapping events changed in the same direction or opposite) with an RBP KD profile. Ranking of RBPs based on RBP differential gene expression alone identified 6 out of the 14 splice-ICGS signature (43%) and one of the 6 splicing factor mutation signatures, among the top 5 predictions for each. RRE subtype enrichment analyses (HOMER) detected 4 of the 11 splice-ICGS signatures (36%, the remaining three signatures did not have a KD profile associated for any of the top 5 RBP) and one of the 6 splicing factor mutation signatures. The CLIP-Seq enrichment (RELI pipeline) alone found concordance for 8 of 14 splicing signatures (57%) and 2 of the 6 splicing factor mutation signatures, however, the specificity of these KD RBP predictions was low (e.g., *SRSF1* was the only KD associated RBP for five signatures of the eight). Hence, the combined logistic regression model substantially improved RBP predictions based on available gold-standards (RBP mutation) or silver-standards (RBP KD concordance), with increased diversity of concordant RBP predictions.

**Section 4**: **Survival Analysis of OncoSplice Subtypes**

To perform survival analysis, we applied the cox proportional hazard test in R using the survival package. The survival analyses were performed for the splice-ICGS subtypes (binary outcomes) or continuous outcomes (e.g., *WDR77* expression) as a multivariate analysis. Several previously reported survival significant variables such as age, cytogenetics, *NPM1* mutations, *FLT3* mutations, white blood cell count, *PML-RARA*, and *MLL* fusions were considered as confounding variables and were included in the model. Clinical annotations were obtained from the TCGA and TARGET initiative websites. No clinical annotations were available for Leucegene samples at the time of submission.

**Section 5: U2AF1 RRE Analysis**

To evaluate differences in *U2AF1* binding site specificity, we used the weblogo software (<http://weblogo.berkeley.edu/logo.cgi>), similar to previously described ^18^. In brief, the nucleotide frequency at the 3’ splice site for each alternative cassette exon (12nt to the -3nt position upstream) was reported for each of the queried splicing signatures.

**Section 6: Splicing Functional Prediction Analysis**

The functional impact of reported splicing events was assessed using initial in silico predictions for global results and manually verified for individual reported events, as described below.

6.1. Global prediction results.

To obtain global predictions for the functional impact of alternative splicing events, we relied on the previously described isoform prediction workflow in AltAnalyze ^37^. This workflow compares each pair of compared exon-exon junctions to all possible aligning mRNA isoforms in the Ensembl and UCSC mRNA databases. Where no aligning protein isoform exists for aligning mRNAs, in silico translation is performed using the Bio-Python library (i.e., the longest translation frame that begins with a UniProt N-terminal peptide sequence). Since different mRNA isoforms aligning to the same junction can encode proteins of different lengths, this workflow takes both the complete exon structure and protein coding potential into account to ensure predictions are maximally conservative. Specifically, AltAnalyze contains an isoform prioritization function (<https://github.com/nsalomonis/altanalyze/blob/master/build_scripts/IdentifyAltIsoforms.py>) that iterates through isoforms aligning to the inclusion-junction (I_incl_) and isoforms aligning to exclusion-junction (I_excl_), and compares each to determine: 1) the fraction of amino acids differing between I_incl_ and I_excl_, 2) the number of amino acids differing and 3) the number of common exons in I_incl_ and I_excl_, in the listed order of priority. Those isoform pairs with the lowest fraction of differing residues are selected, with priority given to Ensembl annotated isoforms. The resulting isoform pairs are compared to evaluate differences in: 1) overall protein length, 2) annotated domain composition (InterPro and UniProt - peptide sequence search) and 3) Ensembl annotated nonsense mediated decay (NMD) annotations.

6.2 Confirmation of protein predictions.

For examples evaluated in this manuscript (**Fig. 3i-k**), we performed a secondary evaluation of protein domain and coding potential. First, splicing events were confirmed among patients using SashimiPlot visualization in IGV (**ED Fig. 3**, **4**). Second, included or excluded exons that specifically aligned to an Ensembl annotated NMD isoform or truncated isoform (<50% protein length of the alternative isoform) were determined based on visualization of the exon aligning sequence in the UCSC genome browser with the isoform track. In cases where no Ensembl isoforms exist, the alternatively spliced exon or intron sequences were selected from the UCSC Genome Browser DAS server and inserted or removed, respectively, from the closest matching mRNA isoform. These isoforms then were then subjected to in silico translation using the ExPasy translation service (<https://web.expasy.org/translate>) to derive protein-level predictions.

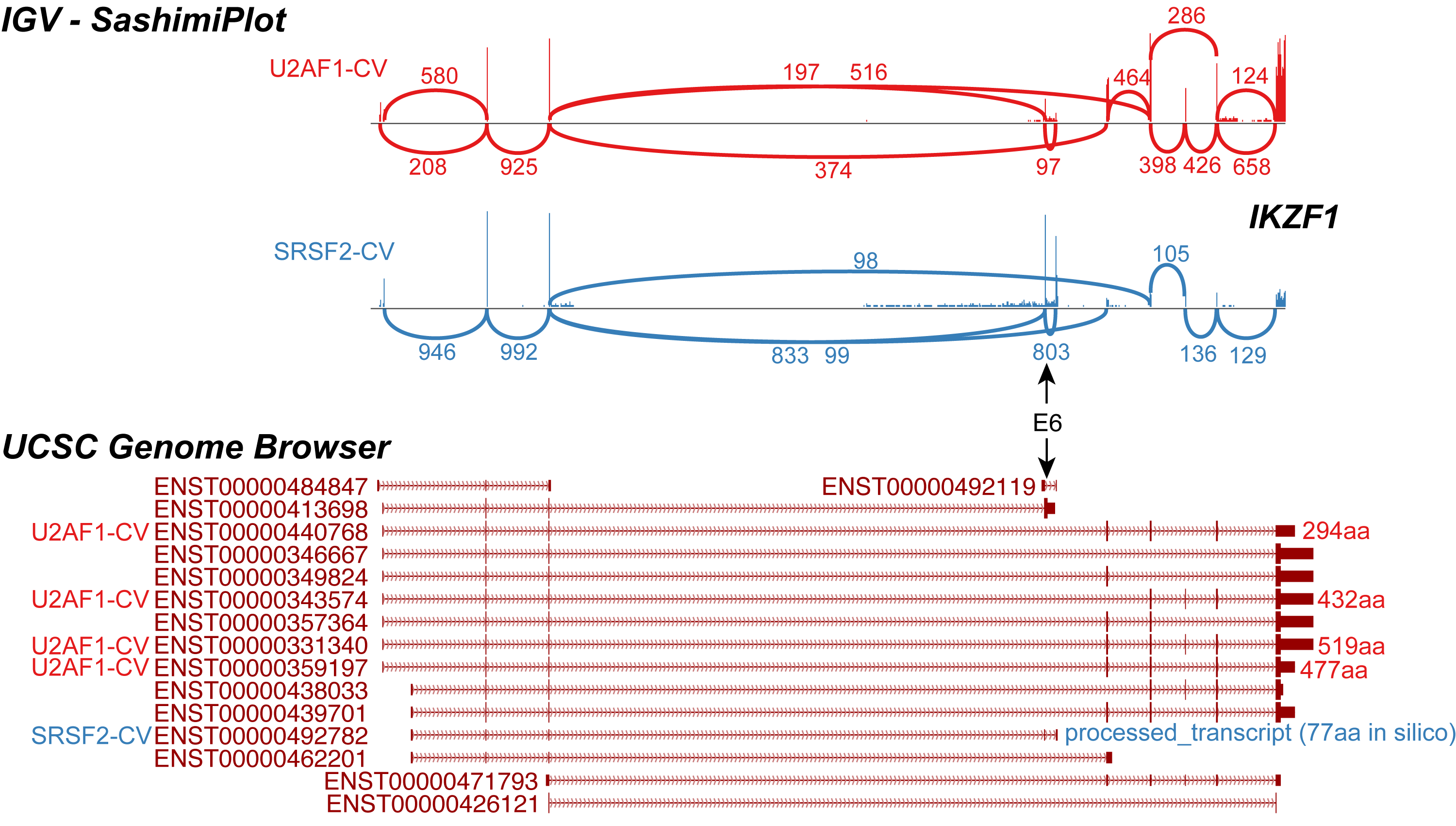

Confirmation of alternative splicing for Ensembl aligning junctions. Decreased inclusion of IKZF1 exon 6 (E6) was identified in *U2AF1*-LIKE versus *SRSF2*-LIKE patients. Associated possible isoforms based on the most frequently expressed exon-exon junctions (IGV, SashimiPlot view) compared to known Ensembl isoforms. Annotated protein isoform lengths are shown to the right of all possible transcripts associated with *U2AF1*-LIKE and *SRSF2*-LIKE samples.

**Section 7: Application of OncoSplice to Leucegene**

To identify both novel and previously implicated splicing subtypes in adult AML, we applied the OncoSplice workflow to Leucegene RNA-Seq data. We remained blinded to the genetics of these patients, the majority of which have not been previously described in the literature. This dataset is composed of primary Bone Marrow or blood samples collected at the time of AML diagnosis. Our analyses were applied only to 367 out of the 438 samples, which do not include mutational or clinical metadata (GSE62190, GSE66917, GSE67039). The remaining samples, which possess mutational information (MISTIQ database), were reserved for supervised classification and subtype validation (GSE49642, GSE52656). Prior to running the analysis, we excluded one sample from the Leucegene dataset (SRS876149) with >50% missing values relative to all other samples. The results from each step of this analysis are shown in **ED** **Fig. 1j**. The following steps were automatically applied by splice-ICGS.

a) Splice-ICGS Round 1 analysis. Splice-ICGS was run using its default options (rho=0.4, cosine HOPACH clustering) on the 125,388 splicing events detected by MultiPath-PSI. Splice-ICGS excluded splicing events with > 25% missing values in all samples, reducing the dataset from ~125K to ~88K splicing events. In its first round, splice-ICGS identified 2 major splicing clusters, *U2AF1*-LIKE and *SRSF2*-LIKE. Stringent and expanded versions of these signatures were reported for these two subtypes as described in Section 1.3.2.
b) Correlation Depletion. Following sample cluster identification, the NMF resultant signatures (W matrix) were correlated to all the splicing events and events correlated to these two signatures were removed. Approximately, 43,000 splicing events remained following this correlation analysis, which were considered for the next iteration of splice-ICGS.
c) Round 2 splice-ICGS and detection of other splicing signatures. 15 additional splicing clusters were identified in the next round, with the software using a rank=30 to identify the sample clusters.
d) Round 2 sample cluster identification. From the 30 splicing patient subtypes, 15 subtypes were excluded by the software based on the two criterions described in Section 1.3.2. This step is necessary to exclude inherently redundant subtypes that will occur due to a larger number of allowed NMF ranks (30). The resulting 15 splicing subtypes were enriched for known RBP mutations (*U2AF1-*S34, *SRSF2-*P95, *SF3B1*) and oncofusions (*MLL* fusions, *RUNX1* fusions, *CBFB* and *PML-RARA*). Mutational enrichment revealed that *NPM1* and *FLT3*-ITD were enriched in two of the subtypes whereas *TP53* mutations were enriched in two other subtypes. We report both stringent and expanded versions of the MLL subtype; however, for downstream analyses, splicing events associated with the genetically defined MLLs were primarily evaluated.
e) Correlation Depletion on Round 2. Splicing events were then again excluded following the second round of splice-ICGS, corresponding to the 15 NMF splicing subtypes, resulting in 32,000 remaining splicing events.
f) Round 3 sample cluster identification. Following splice-ICGS and rank identification, HOPACH clustering identifies only one remaining cluster (rank=1). At this point, the workflow defaults to using k-means to separate the samples into clusters rather than NMF. Performing k-means clustering (k=2) on the Guide 3 splice-ICGS file results in two clusters, with one cluster containing 5 samples and the other cluster containing all remaining samples. Differential splicing analyses results in omitting the larger k-means cluster (n=361 samples) since no significant events are associated with the subtype, whereas the smaller cluster (n=5) had > 100 significant splicing events associated with it. All 5 samples in the resulting final subtype contained *ZRSR2* mutations.

Summary: The final splice-ICGS analysis produced 18 unique splicing subtypes from the Leucegene dataset, with 11 subtypes being enriched in either splicing factor mutations, oncofusions or other AML prior associated variants. The *SRSF2*-P95 signature was split into two signatures based on our subsequent variant analysis (*SRSF2*-P95 and *SRSF2*-8AA deletion). Splicing events associated with each subtype were obtained with MultiPath-PSI using the default δPSI cutoff of >0.1, Benjamini Hochberg adjusted empirical Bayes moderated t-test p<0.05, and requiring detection of values in over 75% of patient samples for each compared sample group. It is further observed that the expanded subtypes for *U2AF1*-LIKE and *SRSF2*-LIKE better match the observed splicing signatures reported by the original Guide-3 and NMF round 1 results and are associated with graded rather than discrete signatures. Both stringent and expanded signatures are supplied online (<https://www.synapse.org/#!Synapse:syn12103684>). Notably in several cases, one or two patients with known splicing factor mutations (*SRSF2*-P95, *SRSF2*-8AA, *SF3B1*, *U2AF1*-S34) were not assigned to the proper patient subtypes by splice-ICGS. Similarly, multiple patients with oncofusions were mis-called (*CBFB*-fusions, *MLL*-fusions, *RUNX1*-fusions) in the above analyses. Although for downstream analyses we revised the subtype associations to only include patients with the confirmed genetic lesions, **ED Table 1** provides both the original and revised annotations.
